# *Fmr1* KO causes delayed rebound spike timing in mediodorsal thalamocortical neurons through regulation of HCN channel activity

**DOI:** 10.1101/2025.02.02.636122

**Authors:** Gregory J. Ordemann, Polina Lyuboslavsky, Alena Kizimenko, Audrey C. Brumback

## Abstract

**Background:** The neurodevelopmental disorder Fragile X syndrome (FXS) results from hypermethylation of the *FMR1* gene which prevents FMRP production. FMRP modulates the expression and function of a wide variety of proteins, including voltage-gated ion channels such as Hyperpolarization-Activated Cyclic Nucleotide gated (HCN) channels, which are integral to rhythmic activity in thalamic structures. Thalamocortical pathology, particularly involving the mediodorsal thalamus (MD), has been implicated in neurodevelopmental disorders. MD connectivity with mPFC is integral to executive functions like working memory and social behaviors that are disrupted in FXS.

**Methods:** We used a combination of retrograde labeling and *ex vivo* brain slice whole cell electrophysiology in 40 wild type and 42 *Fmr1* KO male mice to investigate how a lack of *Fmr1* affects intrinsic cellular properties in lateral (MD-L) and medial (MD-M) MD neurons that project to the medial prefrontal cortex (MD→mPFC neurons).

**Results:** In MD-L neurons, *Fmr1* knockout caused a decrease in HCN-mediated membrane properties such as voltage sag and membrane afterhyperpolarization. These changes in subthreshold properties were accompanied by changes in suprathreshold neuron properties such as the variability of action potential burst timing.

**Conclusions:** In *Fmr1* knockout mice, reduced HCN channel activity in MD→mPFC neurons impairs both the timing and magnitude of HCN-mediated membrane potential regulation. Changes in response timing may adversely affect rhythm propagation in *Fmr1* KO thalamocortical circuitry. MD thalamic neurons are critical for maintaining rhythmic activity involved in cognitive and affective functions. Understanding specific mechanisms of thalamocortical circuit activity may lead to therapeutic interventions for individuals with FXS.

## Introduction

Fragile X syndrome (FXS) is a neurodevelopmental condition characterized by cognitive and social / emotional challenges that partially localize to the prefrontal brain network, which includes mediodorsal thalamus (MD) and prefrontal cortex (PFC). MD contributes to executive functioning and social motivation including attention, working memory, behavioral flexibility, cognitive flexibility, and social motivation (1–10). Changes in thalamocortical connectivity are implicated in autism spectrum disorders (11,12). Studies of executive dysfunction in FXS have identified deficits in working memory in human subjects (13) and cognitive flexibility in a mouse model of FXS (14–16). Previous work has demonstrated that the intrinsic excitability of the neurons that project from the prefrontal cortex to thalamus is disrupted in a mouse model of FXS (17,18). However, the the physiology of the neurons in the mediodorsal (MD) thalamus that provide ascending input to the prefrontal cortex are altered by a lack of *Fmr1* remains unknown.

The two major projections from MD to medial PFC (MD→mPFC) arise from the medial (MD-M) and lateral (MD-L) subnuclei (10,19). Previously, we found that MD-L→mPFC neurons have more Hyperpolarization-Activated Cyclic Nucleotide-Gated (HCN) channel activity, shorter membrane time constants, lower membrane resistance, and require stronger current injections to generate action potentials compared to MD-M→mPFC neurons (19). Animal models of FXS show that intrinsic ion channelopathies can result in changes to synaptic processing and plasticity (20–23). Additionally, HCN channelopathies have been identified in multiple brain regions in mouse models of FXS (18,20,24). The importance of the prefrontal thalamocortical network to executive and social / emotional function suggests MD involvement in cognitive and behavioral symptoms observed in patients with FXS. We hypothesized that *Fmr1* KO MD→mPFC neurons would have intrinsic and circuit dysfunction, in part, due to ion channel dysfunction. To test this hypothesis, we used a retrograde tracer to fluorescently label MD neurons that project to prelimbic and infralimbic cortices in mouse (MD→mPFC neurons). In *ex vivo* thalamic slices from adult mice, we used whole cell current-clamp recordings to measure subthreshold and suprathreshold physiological properties of labeled MD-M→mPFC and MD-L→mPFC neurons. We compared *Fmr1* knockout (KO) animals to their wildtype (WT) littermates. Loss of *Fmr1* caused MD-L→mPFC neurons to display less HCN channel activity, which caused a mild increase in input resistance (*R_N_*). Loss of *Fmr1* did not cause major changes in action potential generation in response to direct current injections. However, MD-L→mPFC neurons in *Fmr1* KO mice showed dampening of HCN-dependent membrane resonator dynamics. First, in *Fmr1* KO mice, MD-L→mPFC neurons had lower amplitude post-depolarization afterhyperpolarizations (AHP). Second, these same neurons showed a slowing of rebound spiking following release of hyperpolarization. These differences in cellular physiology related to membrane resonance have implications for slow wave oscillations in thalamic neurons which may influence prefrontal network activity in FXS with broad implications for sleep and cognitive function.

## Methods and Materials

### Animals

All experiments were conducted in accordance with procedures established by the Institutional Animal Care and Use Committee (IACUC) at The University of Texas. Male mice (8-12 weeks) were used due to the significantly greater prevalence of FXS in males. *Fmr1* het females (gift of Kimberly Huber, UT Southwestern) were crossed with wildtype C57Bl/6J males from Jackson Labs (stock #000664) to yield *Fmr1^+/y^* (wildtype) and *Fmr1* KO mice that were group-housed with same sex littermates in open-top cages in reverse lighting conditions (9am-9pm dark) and fed *ad libitum*.

### Fluorescent labeling of specific neuron populations

Fluorescent labeling of wildtype and *Fmr1* KO MD-L→mPFC and MD-M→mPFC neurons with cholera toxin subunit B (CTB, 500 µg / 100 µL, Molecular Probes, Thermo Fisher Scientific), was performed as in (19) . mPFC injection (450 nL at 100 nL/minute) was delivered to coordinates (in mm relative to Bregma): –1.7 anterior– posterior (AP), +0.3 mediolateral (ML) and –2.75 dorsoventral (DV). After 3-6 days brain tissue was processed for histology or electrophysiological recording. Experiments were performed after 3-6 days on neurons located in MD from bregma = -1.00 to bregma = -1.50. Neurons recorded anterior to this range were considered part of a transition area in MD and excluded from analysis (25). In this manuscript WT and *Fmr1* KO refers to “WT MD-L” and “*Fmr1* KO MD-L” unless otherwise indicated.

### Histology

Injected animals used for histological confirmation of injection location were transcardially perfused with paraformaldehyde (PFA, Sigma-Aldrich) 4% in phosphate-buffered saline (1x PBS). Tissue was sectioned in 50 µm thick coronal slices with DAPI-containing mounting medium (VectaShield HardSet with DAPI, Vector laboratories) and slices containing mPFC and MD were imaged at 5x using Zeiss Axio Imager 2.

### Slice preparation

250 μm thick slices (Leica VT1200) were prepared from mice 8 to 12 weeks old after intraperitoneal injection of ketamine / xylazine (90 / 10 mg/kg; Acor/Dechra). Mice were perfused with cutting solution containing (in mM): 205 sucrose, 25 NaHCO_3_, 2.5 KCl, 1.25 NaH_2_PO_4_, 7 MgCl_2_, 7 dextrose, 3 Na pyruvate, 1.3 sodium ascorbate, and 0.5 CaCl_2_ bubbled with 95% O_2_ / 5% CO_2_. Slices were incubated in holding solution containing (in mM): 125 NaCl, 25 NaHCO_3_, 2.5 KCl, 1.25 NaH_2_PO_4_, 25 dextrose, 2 CaCl_2_, 2 MgCl_2_, 1.3 sodium ascorbate, and 3 Na pyruvate at 37±1°C for 30 minutes then kept for at least 30 minutes at room temperature before recording began.

### Intracellular recordings

Artificial cerebrospinal fluid (ACSF) contained (in mM): 125 NaCl, 25 NaHCO_3_, 12.5 dextrose, 2.5 KCl, 1.25 NaH_2_PO_4_ 2 CaCl_2_ and 1 MgCl_2_. Slices were continuously perfused with ACSF in an immersion chamber (Warner Instruments) with temperature maintained at 32.5 ± 1°C (Warner Instruments TC-324C). We did not add synaptic blockers to the ACSF unless otherwise specified. In a subset of experiments ZD7288 (20 μM; Tocris Cat#1000) or TTX (0.5 μM; Abcam Cat# were added to extracellular ACSF.

Somatic whole-cell patch recordings were obtained from retrogradely-labeled neurons in the medial (MD-M) or lateral (MD-L) subnuclei of WT and *Fmr1* KO mice, using DODT (Zen 2.5 blue addition, Zen pro) contrast microscopy and epifluorescence on an upright microscope (Zeiss Examiner D1). Patch electrodes (tip resistance = 3–6 MΩ) were filled with the following (in mM): 118 K-gluconate, 10 KCl, 10 HEPES, 4 MgATP, 1 EGTA, 0.3 Na3GTP and 0.3% biocytin (pH adjusted to 7.2 with KOH; 282 mOsM). For some cells, 16 µM Alexa 488 or Alexa 594 was added to the internal solution to visualize the dendritic arbor under epifluorescence. Recordings were made with Clampex 10.7 software running a Multiclamp 700B (Molecular Devices). Signals were digitized at 20 kHz and lowpass filtered at 4 kHz.

Data were collected at resting membrane potential (RMP) and -65 ± 3 mV. Unless stated otherwise, all data reported here were taken from recordings performed at -65 mV. Experiments were discontinued if series resistance rose above 30 MΩ or action potentials failed to overshoot 0 mV. Liquid junction potential was estimated to be 14.3 mV using Patchers Power Tools (IGORpro 7, Wavemetrics). Liquid junction potential was not corrected. The data sets were sampled from 40 WT and 42 *Fmr1* KO mice.

### Electrophysiological properties

The analysis of electrophysiological data was performed as in (19) . Briefly, RMP, τ, and *R_N_*. were measured from voltage responses to subthreshold current injections. Analyses were performed on a series of current injections ranging from -60 to 60 pA in 5 pA intervals for 1000 ms or from -250 to 350 in 25 pA intervals. Voltage sag is the difference between peak hyperpolarization and steady state voltage deflection in the trace with a peak hyperpolarization of -100 mV. Afterhyperpolarzation is the difference between peak hyperpolarization and baseline membrane potential following the offset of the first current stimulus to have 12 or more action potentials. In **Figs. 3F & I** and **Fig. 5C**, neurons in the wildtype ZD wash-on experiments were previously analyzed in (19).

### Burst and tonic action potentials

We measured action potential threshold as the point at which the third derivative of the membrane potential exceeded 0.3 V/s^3^. Firing frequency measurements for tonic firing included sweeps in which no action potentials were fired, however, frequency measurements for burst action potentials excluded instances in which no action potentials were present. Accommodation was measured as the slope of the linear relationship between the interspike intervals for each successive action potential during the first current step to elicit ≥12 action potentials. Low threshold calcium spikes (LTS) were isolated with 0.5 μM TTX. LTS analysis was performed on the first depolarizing step that elicited an LTS or on the hyperpolarizing step that reached -100 mV for rebound LTSs. *α*EPSPs were generate using the equation 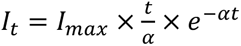, where 𝐼_t_is the current 𝐼 at elapsed time 𝑡. 𝐼_max_ is the maximum current. 𝛼 is a constant representing the decay rate.

### Statistics

We used the ‘sampsizepwr’ function in MATLAB to calculate sample sizes based on preliminary data. To detect a difference in membrane time constant of 25% with a standard deviation of 20 ms, given α of 0.05 and power of 0.8, we estimated at least 16 cells per group. All quantifications are presented in Tables 1-2. Data compared with a two-way ANOVA or Fixed effects (type III) are reported as mean ± 95% CI. Data compared with Mann-Whitney or Wilcoxon tests are presented with all data points and median with 95% confidence intervals. We defined ɑ as p < 0.05. Effect size is expressed as *η*^2^ and only calculated only when p<0.05. *η*^2^ effect sizes are defined as small – 0.01, medium – 0.06, and large – 0.14 (26). Quantifications were performed using custom-written code in MATLAB 2022b (Mathworks). Statistical analyses were performed using Prism version 8.0.0 (GraphPad). Graphs were made using GraphPad Prism and figures were made using Adobe Illustrator version 24.3.

**Table 1:** Descriptive statistics and test statistics of pairwise comparisons.

| Comparison | groups | n-value<br>(cells/animals) | Mean ± SEM | median [-/+ 95% CI] | Statistical<br>test | Test<br>Statistic | p-value | eta-<br>squared | Median<br>difference<br>[-/+ 95% CI] |
| --- | --- | --- | --- | --- | --- | --- | --- | --- | --- |
| Resting $V_m$ (mV) | MD-L | 39/29 | -60.92 ± 1.1 | -62 [-65/-58] | Mann-Whitney | U = 1129 | 0.87 | | -1 [-2/2] |
| | MD-L *Fmr1* KO | 59/38 | -60.94 ± 0.89 | -63 [-65/-61] | | | | | |
| Resting $V_m$ (mV) | MD-M | 20/14 | -62.2 ± 1.89 | -63.5 [-69/-61] | Mann-Whitney | U = 237 | 0.09 | | 3.5 [-1/8] |
| | MD-M *Fmr1* KO | 33/21 | -58.79 ± 1.42 | -60 [-63/059] | | | | | |
| Tau (ms) | MD-L | 38/27 | 58.01 ± 3.76 | 52.81 [46.51/63.34] | Mann-Whitney | U = 1070 | 0.92 | | 1.45<br>[-6.96/6.58] |
| | MD-L *Fmr1* KO | 57/36 | 55.69 ± 2.19 | 54.27 [50.31/57.68] | | | | | |
| Tau (ms) | MD-M | 20/13 | 68.35 ± 4.25 | 66.85 [60.25/78.31] | Mann-Whitney | U = 310 | 0.86 | | 0.47<br>[-10.59/12.32] |
| | MD-M *Fmr1* KO | 32/19 | 69.36 ± 3.86 | 67.32 [59.18/80.89] | | | | | |
| $R_N$ (MΩ) | MD-L | 36/28 | 253.1 ± 19.41 | 216 [180.2/289.3] | Mann-Whitney | U = 633.5 | 0.037 | 0.052 | 82.95<br>[3.23/110.3] |
| | MD-L *Fmr1* KO | 48/33 | 308.8 ± 18.45 | 298.9 [243.7/345.6] | | | | | |
| $R_N$ (MΩ) | MD-M | 20/13 | 382.1 ± 26.34 | 397.2 [311.7/487.3] | Mann-Whitney | U = 190 | 0.087 | | 75.45<br>[-16.8/172.2] |
| | MD-M *Fmr1* KO | 27/17 | 462.8 ± 21.36 | 472.6 [331.8/610.4] | | | | | |
| Voltage sag at<br>-100 mV (mV) | MD-L | 40/30 | -10.27 ± 0.77 | -10.07 [-12.89/-7.96] | Mann-Whitney | U = 791 | 0.004 | 0.08 | 3.19 [1.03/4.95] |
| | MD-L *Fmr1* KO | 60/39 | -7.48 ± 0.54 | -6.88 [-8.66/-5.84] | | | | | |
| Voltage sag at<br>-100 mV (mV) | MD-M | 18/12 | -5.45 ± 0.94 | -4.52 [-7.5/-2.94] | Mann-Whitney | U = 237 | 0.24 | | 1.71<br>[-0.66/2.88] |
| | MD-M *Fmr1* KO | 33/21 | -3.95 ± 0.45 | -2.81 [-5.41/-2.14] | | | | | |
| $R_N$ (MΩ) - ZD<br>wash-on | MD-L pre | 12/7 | 382.5 ± 60.2 | 405 [190/480] | Wilcoxon | W = 73 | 0.002 | 3.53 | 215 [70/280] |
| | MD-L post | 12/7 | 578.3 ± 60.04 | 555 [380/720] | | | | | |
| $R_N$ (MΩ) - ZD<br>wash-on | MD-L *Fmr1* KO pre | 10/6 | 430.0 ± 44.02 | 445 [240/550] | Wilcoxon | W = 55 | 0.002 | 3.14 | 90 [30/270] |
| | MD-L *Fmr1* KO post | 10/6 | 560 ± 47.89 | 575 [460/670] | | | | | |
| Voltage sag at -100<br>mV (mV) - ZD<br>wash-on | MD-L pre | 12/7 | -9.41 ± 1.01 | -8.82 [-11.34/-7.04] | Wilcoxon | W = 78 | 0.0005 | 3.12 | 8.74 [7.15/10.93] |
| | MD-L post | 12/7 | -0.34 ± 0.13 | -0.16 [-0.69/-0.03] | | | | | |
| Voltage sag at -100<br>mV (mV) - ZD<br>wash-on | MD-L *Fmr1* KO pre | 9/6 | -4.77 ± 0.90 | -5.30 [-7.74/-1.43] | Wilcoxon | W = 45 | 0.004 | 3.16 | 5.59 [0.82/7.64] |
| | MD-L *Fmr1* KO post | 9/6 | -0.26 ± 0.09 | -0.11 [-.61/-0.04] | | | | | |
| Afterhyperpolarization<br>(mV) | MD-L | 41/31 | -11.96 ± 0.92 | -11.7 [-15.43/-9.24] | Mann-Whitney | U = 828 | 0.036 | 0.05 | 2.17 [0.2/5.54] |
| | MD-L *Fmr1* KO | 54/37 | -9.14 ± 0.8 | -9.52 [-11.03/-6.39] | | | | | |
| Afterhyperpolarization<br>(mV) - ZD wash-on | MD-L pre | 11/8 | -13.5 ± 1.41 | -13.73 [-18.65/-6.45] | Wilcoxon | W = 66 | 0.001 | 3.13 | 16.21<br>[11.94/24.63] |
| | MD-L post | 11/8 | 4.1 ± 1.12 | 5.35 [-0.82/7.32] | | | | | |
| Afterhyperpolarization<br>(mV) - ZD wash-on | MD-L *Fmr1* KO pre | 9/6 | -9.08 ± 2.08 | -8.65 [-17.81/-2.51] | Wilcoxon | W = 45 | 0.004 | 3.16 | 10.4 [0.52/17.78] |
| | MD-L *Fmr1* KO post | 9/6 | 0.34 ± 0.61 | -0.06 [-1.21/1.86] | | | | | |
| Rebound spike timing | MD-L | 39/29 | 33.29 ± 2.92 | 29.2 [24.25/31.4] | Mann-Whitney | 619.5 | 0.0014 | 0.13 | 10.03<br>[3.75/16.54] |
| | MD-L *Fmr1* KO | 52/36 | 45.6 ± 3.26 | 39.23 [33.5/46.5] | | | | | |
| Rebound spike timing<br>- ZD washon | MD-L pre | 7/5 | 38.41 ± 8.84 | 34.25 [19.15/89.45] | Wilcoxon | W = 28 | 0.016 | 3.2 | 68.25<br>[31.85/104.4] |
| | MD-L post | 7/5 | 107.5 ± 8.91 | 110.7 [76.2/139.1] | | | | | |
| Rebound spike timing<br>- ZD washon | MD-L *Fmr1* KO pre | 6/4 | 67.97 ± 12.12 | 60.73 [33/116.2] | Wilcoxon | W = 7 | 0.56 | | 11.95<br>[-31.7/98.05] |
| | MD-L *Fmr1* KO post | 6/4 | 89.13 ± 16.22 | 84.68 [34.15/156.3] | | | | | |
| Accommodation Index -<br>Tonic firing: RMP | MD-L | 9/9 | 0.002 ± 0.0007 | 0.0016 [0.00035/0.0037] | Mann-Whitney | U = 140.5 | 0.3 | | 0.0009<br>[-0.001/0.003] |
| | MD-L *Fmr1* KO | 5/5 | 0.003 ± 0.0006 | 0.0026 [0.0014/0.0051] | | | | | |
| Action potential<br>threshold - Tonic<br>firing: RMP (mV) | MD-L | 9/9 | -32.82 ± 0.99 | -33.07 [-36.25/-30.52] | Mann-Whitney | U = 19 | 0.7 | | -2.8<br>[-7.24/6.09] |
| | MD-L *Fmr1* KO | 5/5 | -33.63 ± 2.4 | -35.87 [-38.28/-26.98] | | | | | |
| Time to first spike -<br>Tonic firing: RMP<br>(ms) | MD-L | 9/9 | 216 ± 81.43 | 147 [11/578] | Mann-Whitney | U = 19 | 0.94 | | -24 [-414/153] |
| | MD-L *Fmr1* KO | 5/5 | 214.2 ± **103.6** | 123 [50/622] | | | | | |
| Accommodation Index -<br>Tonic firing:<br>Mibefradil | MD-L | 8/6 | 0.0045 ± 0.001 | 0.004 [0.00047/0.014] | Mann-Whitney | U = 12 | 0.28 | | -0.0011<br>[-0.005/0.001] |
| | MD-L *Fmr1* KO | 5/4 | 0.0024 ± 0.0006 | 0.002 [0.0009/0.0044] | | | | | |
| Action potential<br>threshold - Tonic<br>firing: Mibefradil (mV) | MD-L | 8/6 | -37.07 ± 1.61 | -37.66 [-42.27/-27.59] | Mann-Whitney | U = 9.5 | 0.14 | | 4.46<br>[-1.56/8.03] |
| | MD-L *Fmr1* KO | 5/4 | -33.98 ± 1.26 | -33.2 [-37.57/-31.03] | | | | | |
| Time to first spike -<br>Tonic firing:<br>Mibefradil (ms) | MD-L | 8/6 | 114 ± 22.58 | 99 [47/190] | Mann-Whitney | U = 14 | 0.93 | | -1 [-92/176] |
| | MD-L *Fmr1* KO | 5/4 | 136 ± 44.24 | 98 [36/292] | | | | | |
| Burst threshold (mV) | MD-L | 30/22 | -38.99 ± 0.65 | -39.69 [-40.77/-37.75] | Mann-Whitney | U = 550 | 0.81 | | 1.17 [-1.67/2.16] |
| | MD-L *Fmr1* KO | 38/28 | -39.02 ± 0.75 | -38.52 [-40.74/-37.66] | | | | | |
| Step burst LTS spike<br>threshold (mV) | MD-L | 9/5 | -48.14 ± 0.97 | -48.43 [-50.84/-45.56] | Mann-Whitney | U = 24 | 0.47 | | -1.36<br>[-4.34/1.39] |
| | MD-L *Fmr1* KO | 7/5 | -49.59 ± 0.74 | -49.9 [-52.05/-47.15] | | | | | |
| Step burst LTS peak time (ms) | MD-L | 9/5 | 87.67 ± 6.81 | 89 [68/110] | Mann-Whitney | U = 24 | 0.47 |  | 13 [-25/44] |
|  | MD-L *Fmr1* KO | 7/5 | 98.71 ± 11.76 | 102 [58/136] |  |  |  |  |  |
| Step burst LTS threshold time (ms) | MD-L | 9/5 | 62.78 ± 7.6 | 64 [41/90] | Mann-Whitney | U = 21.5 | 0.31 |  | 5 [-21/54] |
|  | MD-L *Fmr1* KO | 7/5 | 78.14 ± 11.51 | 69 [38/118] |  |  |  |  |  |
| Rebound AP threshold (mV) | MD-L | 33/24 | -41.07 ± 0.53 | -41.75 [-42.88/-40.56] | Mann-Whitney | U = 607.5 | 0.37 |  | 1.8 [-0.92/2.63] |
|  | MD-L *Fmr1* KO | 42/31 | -40.16 ± 0.75 | -39.95 [-41.9/-38.6] |  |  |  |  |  |
| Rebound LTS spike threshold (mV) | MD-L | 12/7 | -56.54 ± 0.76 | -56.39 [-59.45/-54.66] | Mann-Whitney | U = 57 | 0.87 |  | -0.71 [-2.4/2.45] |
|  | MD-L *Fmr1* KO | 10/5 | -56.54 ± 0.69 | -57.09 [-58.57/-53.6] |  |  |  |  |  |
| Rebound LTS peak time (ms) | MD-L | 12/7 | 50.58 ± 6.08 | 48.5 [30/69] | Mann-Whitney | U = 40 | 0.2 |  | 9.5 [-6/37] |
|  | MD-L *Fmr1* KO | 10/5 | 64.8 ± 7.9 | 58 [41/90] |  |  |  |  |  |
| Rebound LTS threshold time (ms) | MD-L | 12/7 | 45.58 ± 7.43 | 43.5 [24/59] | Mann-Whitney | U = 27 | 0.028 | 0.22 | 15.5 [1/68] |
|  | MD-L *Fmr1* KO | 10/5 | 77.4 ± 11.4 | 59 [47/127] |  |  |  |  |  |
| Accommodation Index - Tonic firing: MD-M | MD-M | 4/4 | 0.0018 ± 0.00071 | 0.002 [0.00004/0.0029] | Mann-Whitney | U = 8 | 0.73 |  | -0.0008 [-0.003/0.004] |
|  | MD-M *Fmr1* KO | 5/5 | 0.0016 ± 0.00094 | 0.0013 [-0.00022/0.0049] |  |  |  |  |  |
| Action potential threshold - Tonic firing: MD-M (mV) | MD-M | 4/4 | -32.58 ± 2.76 | -33.1 [-37.41/-26.7] | Mann-Whitney | U = 7 | 0.56 |  | 2.19 [07.23/15.18] |
|  | MD-M *Fmr1* KO | 5/5 | -29.85 ± 2.46 | -30.91 [-36.27/-21.98] |  |  |  |  |  |
| Time to first spike - Tonic firing: MD-M (ms) | MD-M | 4/4 | 196.3 ± 62.33 | 187 [57/354] | Mann-Whitney | U = 9 | 0.9 |  | -68 [-248/468] |
|  | MD-M *Fmr1* KO | 5/5 | 243.4 ± 106.8 | 119 [37/622] |  |  |  |  |  |
| Burst action potential threshold (mV) | MD-M | 16/11 | -36.09 ± 1.07 | -36.88 [-38.27/-35.25] | Mann-Whitney | U = 140.5 | 0.16 |  | -1.57 [-3.53/0.63] |
|  | MD-M *Fmr1* KO | 24/14 | -38.28 ± 0.84 | -38.45 [-40.23/-35.06] |  |  |  |  |  |
| Rebound action potential threshold (mV) | MD-M | 18/12 | -37.51 ± 0.83 | -38.5 [-39.42/-36.57] | Mann-Whitney | U = 185 | 0.18 |  | -1.02 [-3.22/0.61] |
|  | MD-M *Fmr1* KO | 27/17 | -39.11 ± 0.61 | -39.52 [-40.99/-37.67] |  |  |  |  |  |
| First αEPSP amplitude | MD-L | 12/10 | 1.85 ± 0.22 | 1.9 [1.13/2.11] | Mann-Whitney | U = 52 | 0.62 | | -0.12 [-0.42/0.73] |
|  | MD-L *Fmr1* KO | 10/10 | 1.94 ± 0.17 | 1.78 [1.46/2.7] |  |  |  |  |  |

**Table 2:**
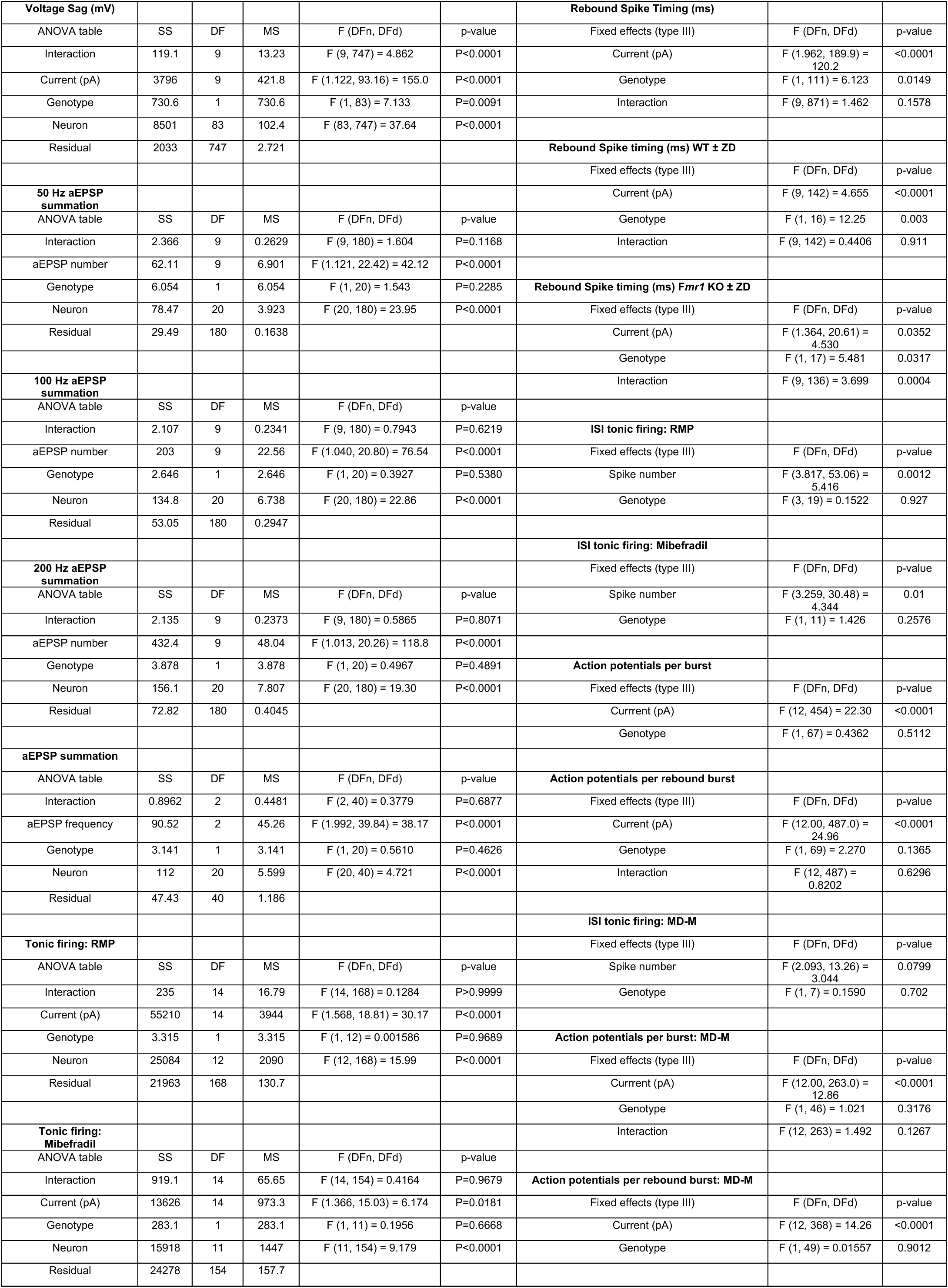
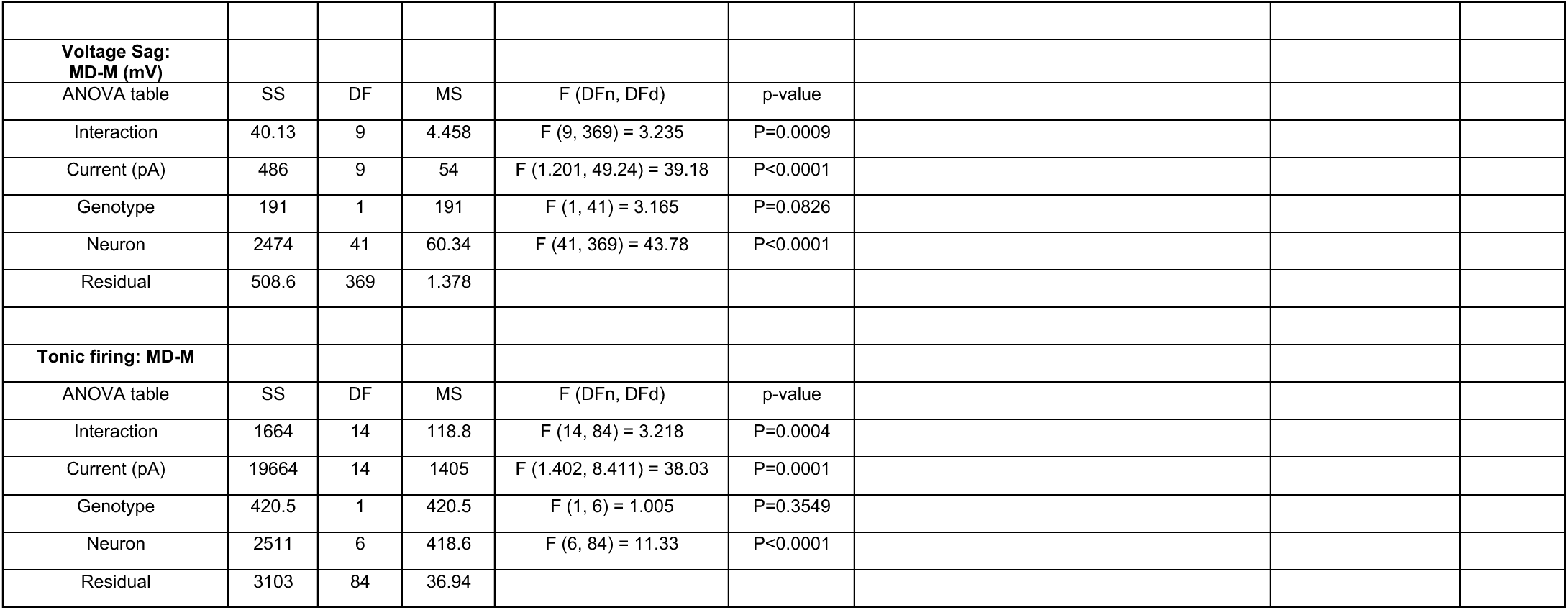
ANOVA and Fixed effects statistics tables for experiments with multiple measurement points.

|  |  |  |  |  |  |  |  |  |
| --- | --- | --- | --- | --- | --- | --- | --- | --- |
| **Voltage Sag (mV)** |  |  |  |  |  | **Rebound Spike Timing (ms)** |  |  |
| ANOVA table | SS | DF | MS | F (DFn, DFd) | p-value | Fixed effects (type III) | F (DFn, DFd) | p-value |
| Interaction | 119.1 | 9 | 13.23 | F (9, 747) = 4.862 | P<0.0001 | Current (pA) | F (1.962, 189.9) = 120.2 | <0.0001 |
| Current (pA) | 3796 | 9 | 421.8 | F (1.122, 93.16) = 155.0 | P<0.0001 | Genotype | F (1, 111) = 6.123 | 0.0149 |
| Genotype | 730.6 | 1 | 730.6 | F (1, 83) = 7.133 | P=0.0091 | Interaction | F (9, 871) = 1.462 | 0.1578 |
| Neuron | 8501 | 83 | 102.4 | F (83, 747) = 37.64 | P<0.0001 |  |  |  |
| Residual | 2033 | 747 | 2.721 |  |  | **Rebound Spike timing (ms) WT ± ZD** |  |  |
|  |  |  |  |  |  | Fixed effects (type III) | F (DFn, DFd) | p-value |
| **50 Hz aEPSP summation** |  |  |  |  |  | Current (pA) | F (9, 142) = 4.655 | <0.0001 |
| ANOVA table | SS | DF | MS | F (DFn, DFd) | p-value | Genotype | F (1, 16) = 12.25 | 0.003 |
| Interaction | 2.366 | 9 | 0.2629 | F (9, 180) = 1.604 | P=0.1168 | Interaction | F (9, 142) = 0.4406 | 0.911 |
| aEPSP number | 62.11 | 9 | 6.901 | F (1.121, 22.42) = 42.12 | P<0.0001 |  |  |  |
| Genotype | 6.054 | 1 | 6.054 | F (1, 20) = 1.543 | P=0.2285 | **Rebound Spike timing (ms) *Fmr1* KO ± ZD** |  |  |
| Neuron | 78.47 | 20 | 3.923 | F (20, 180) = 23.95 | P<0.0001 | Fixed effects (type III) | F (DFn, DFd) | p-value |
| Residual | 29.49 | 180 | 0.1638 |  |  | Current (pA) | F (1.364, 20.61) = 4.530 | 0.0352 |
|  |  |  |  |  |  | Genotype | F (1, 17) = 5.481 | 0.0317 |
| **100 Hz aEPSP summation** |  |  |  |  |  | Interaction | F (9, 136) = 3.699 | 0.0004 |
| ANOVA table | SS | DF | MS | F (DFn, DFd) | p-value |  |  |  |
| Interaction | 2.107 | 9 | 0.2341 | F (9, 180) = 0.7943 | P=0.6219 | **ISI tonic firing: RMP** |  |  |
| aEPSP number | 203 | 9 | 22.56 | F (1.040, 20.80) = 76.54 | P<0.0001 | Fixed effects (type III) | F (DFn, DFd) | p-value |
| Genotype | 2.646 | 1 | 2.646 | F (1, 20) = 0.3927 | P=0.5380 | Spike number | F (3.817, 53.06) = 5.416 | 0.0012 |
| Neuron | 134.8 | 20 | 6.738 | F (20, 180) = 22.86 | P<0.0001 | Genotype | F (3, 19) = 0.1522 | 0.927 |
| Residual | 53.05 | 180 | 0.2947 |  |  |  |  |  |
|  |  |  |  |  |  | **ISI tonic firing: Mibefradil** |  |  |
| **200 Hz aEPSP summation** |  |  |  |  |  | Fixed effects (type III) | F (DFn, DFd) | p-value |
| ANOVA table | SS | DF | MS | F (DFn, DFd) | p-value | Spike number | F (3.259, 30.48) = 4.344 | 0.01 |
| Interaction | 2.135 | 9 | 0.2373 | F (9, 180) = 0.5865 | P=0.8071 | Genotype | F (1, 11) = 1.426 | 0.2576 |
| aEPSP number | 432.4 | 9 | 48.04 | F (1.013, 20.26) = 118.8 | P<0.0001 |  |  |  |
| Genotype | 3.878 | 1 | 3.878 | F (1, 20) = 0.4967 | P=0.4891 | **Action potentials per burst** |  |  |
| Neuron | 156.1 | 20 | 7.807 | F (20, 180) = 19.30 | P<0.0001 | Fixed effects (type III) | F (DFn, DFd) | p-value |
| Residual | 72.82 | 180 | 0.4045 |  |  | Current (pA) | F (12, 454) = 22.30 | <0.0001 |
|  |  |  |  |  |  | Genotype | F (1, 67) = 0.4362 | 0.5112 |
| **aEPSP summation** |  |  |  |  |  |  |  |  |
| ANOVA table | SS | DF | MS | F (DFn, DFd) | p-value | **Action potentials per rebound burst** |  |  |
| Interaction | 0.8962 | 2 | 0.4481 | F (2, 40) = 0.3779 | P=0.6877 | Fixed effects (type III) | F (DFn, DFd) | p-value |
| aEPSP frequency | 90.52 | 2 | 45.26 | F (1.992, 39.84) = 38.17 | P<0.0001 | Current (pA) | F (12.00, 487.0) = 24.96 | <0.0001 |
| Genotype | 3.141 | 1 | 3.141 | F (1, 20) = 0.5610 | P=0.4626 | Genotype | F (1, 69) = 2.270 | 0.1365 |
| Neuron | 112 | 20 | 5.599 | F (20, 40) = 4.721 | P<0.0001 | Interaction | F (12, 487) = 0.8202 | 0.6296 |
| Residual | 47.43 | 40 | 1.186 |  |  |  |  |  |
|  |  |  |  |  |  | **ISI tonic firing: MD-M** |  |  |
| **Tonic firing: RMP** |  |  |  |  |  | Fixed effects (type III) | F (DFn, DFd) | p-value |
| ANOVA table | SS | DF | MS | F (DFn, DFd) | p-value | Spike number | F (2.093, 13.26) = 3.044 | 0.0799 |
| Interaction | 235 | 14 | 16.79 | F (14, 168) = 0.1284 | P>0.9999 | Genotype | F (1, 7) = 0.1590 | 0.702 |
| Current (pA) | 55210 | 14 | 3944 | F (1.568, 18.81) = 30.17 | P<0.0001 |  |  |  |
| Genotype | 3.315 | 1 | 3.315 | F (1, 12) = 0.001586 | P=0.9689 | **Action potentials per burst: MD-M** |  |  |
| Neuron | 25084 | 12 | 2090 | F (12, 168) = 15.99 | P<0.0001 | Fixed effects (type III) | F (DFn, DFd) | p-value |
| Residual | 21963 | 168 | 130.7 |  |  | Current (pA) | F (12.00, 263.0) = 12.86 | <0.0001 |
|  |  |  |  |  |  | Genotype | F (1, 46) = 1.021 | 0.3176 |
| **Tonic firing: Mibefradil** |  |  |  |  |  | Interaction | F (12, 263) = 1.492 | 0.1267 |
| ANOVA table | SS | DF | MS | F (DFn, DFd) | p-value |  |  |  |
| Interaction | 919.1 | 14 | 65.65 | F (14, 154) = 0.4164 | P=0.9679 | **Action potentials per rebound burst: MD-M** |  |  |
| Current (pA) | 13626 | 14 | 973.3 | F (1.366, 15.03) = 6.174 | P=0.0181 | Fixed effects (type III) | F (DFn, DFd) | p-value |
| Genotype | 283.1 | 1 | 283.1 | F (1, 11) = 0.1956 | P=0.6668 | Current (pA) | F (12, 368) = 14.26 | <0.0001 |
| Neuron | 15918 | 11 | 1447 | F (11, 154) = 9.179 | P<0.0001 | Genotype | F (1, 49) = 0.01557 | 0.9012 |
| Residual | 24278 | 154 | 157.7 |  |  |  |  |  |

|  |  |  |  |  |  |
| --- | --- | --- | --- | --- | --- |
| **Voltage Sag:**<br>**MD-M (mV)** |  |  |  |  |  |
| ANOVA table | SS | DF | MS | F (DFn, DFd) | p-value |
| Interaction | 40.13 | 9 | 4.458 | F (9, 369) = 3.235 | P=0.0009 |
| Current (pA) | 486 | 9 | 54 | F (1.201, 49.24) = 39.18 | P<0.0001 |
| Genotype | 191 | 1 | 191 | F (1, 41) = 3.165 | P=0.0826 |
| Neuron | 2474 | 41 | 60.34 | F (41, 369) = 43.78 | P<0.0001 |
| Residual | 508.6 | 369 | 1.378 |  |  |
| **Tonic firing: MD-M** |  |  |  |  |  |
| ANOVA table | SS | DF | MS | F (DFn, DFd) | p-value |
| Interaction | 1664 | 14 | 118.8 | F (14, 84) = 3.218 | P=0.0004 |
| Current (pA) | 19664 | 14 | 1405 | F (1.402, 8.411) = 38.03 | P=0.0001 |
| Genotype | 420.5 | 1 | 420.5 | F (1, 6) = 1.005 | P=0.3549 |
| Neuron | 2511 | 6 | 418.6 | F (6, 84) = 11.33 | P<0.0001 |
| Residual | 3103 | 84 | 36.94 |  |  |

## Results

To investigate the intrinsic effects of *Fmr1* KO on MD neurons projecting to mPFC (“MD→mPFC neurons” but for simplicity referred to as “MD neurons” in this manuscript) we made whole cell current clamp recordings of fluorescently labeled thalamocortical neurons in the MD region of thalamus in acute *ex vivo* mouse brain slices (**Fig. 1A**). MD neurons were visually identified and distinguished as medial (MD-M) or lateral (MD-L) based on the distance from the midline and anatomical landmarks (**Fig. 1B-D**). Both MD-M and MD-L neurons in WT and *Fmr1* KO mice were investigated during this study, however, intrinsic properties only differed between WT and *Fmr1* KO MD-L neurons. As a result, MD-L data is presented in the main b ody of this manuscript (**Figs. 1-8**) while all MD-M data is presented in the supplement (**Supp.** Figs. 2-3).

**Figure 1:**
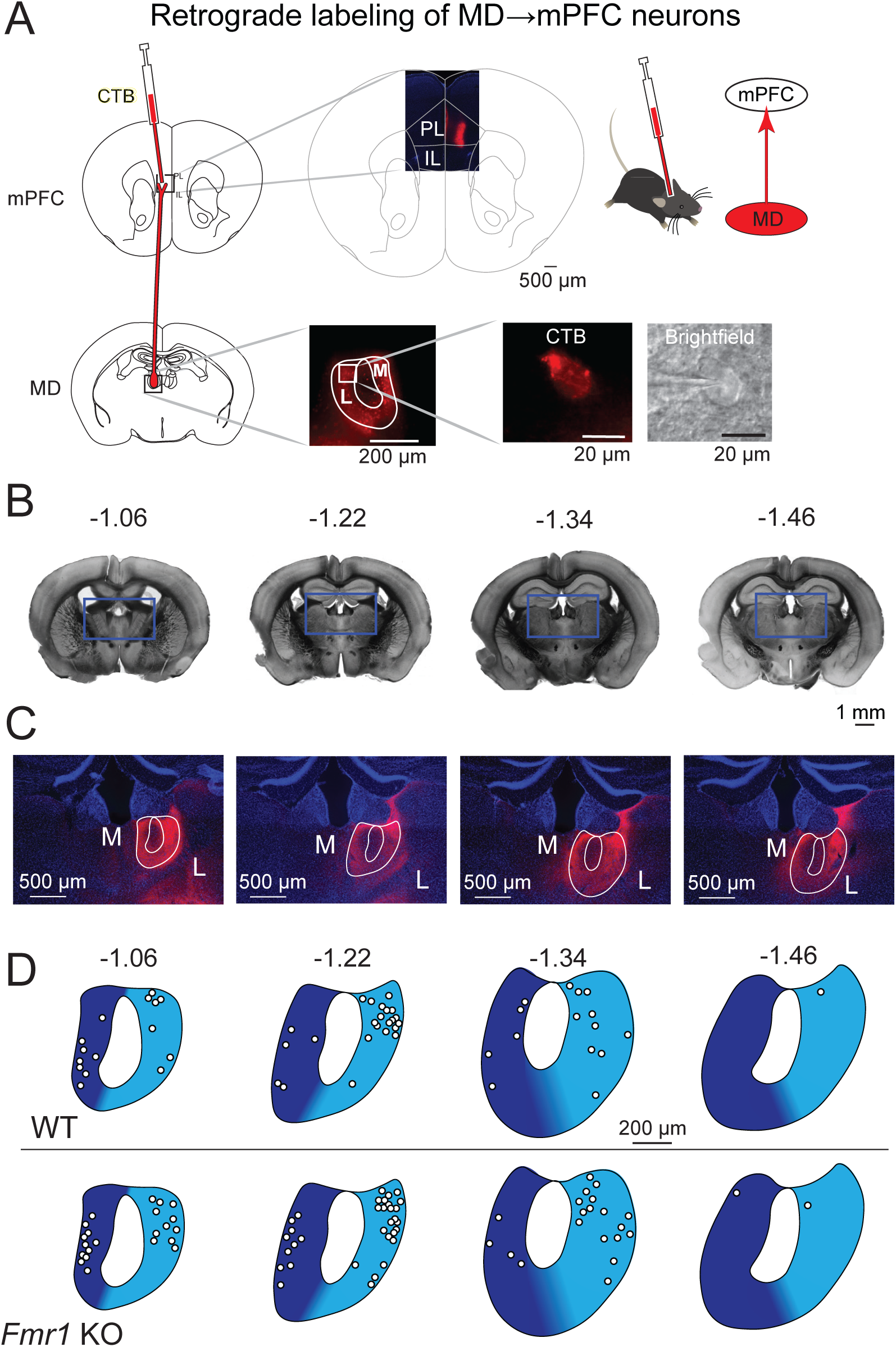
CTB injections in mPFC label MD-L and MD-M neurons that project to mPFC. **A.** Adult wildtype and *Fmr1* KO mice were stereotaxically injected with the retrograde fluorescent tracer CTB into mPFC to target prelimbic and infralimbic cortex. Fluorescently-labeled MD→mPFC neurons were visually identified for patch clamp electrophysiology experiments. **B.** Bright field photomicrographs of coronal mouse brain slices with the estimated distance from Bregma (in mm) where MD→mPFC neurons were patched in this investigation. **C.** Photomicrographs (10x) of the boxed areas from **B** demonstrating MD→mPFC labeled neurons (red) in each of the representative slices. **D.** Map of the approximate locations of recorded neurons reported in this manuscript placed on a single representative atlas drawing for each coronal slice. MD-M is dark blue. MD-L is highlighted in cyan.

### Fmr1 KO MD-L neurons display greater R_N_ compared with WT MD neurons

We measured subthreshold, intrinsic neuronal properties of WT and *Fmr1* KO fluorescently labeled neurons (**Fig. 2A**). We found no difference in either RMP (**Fig. 2B**; Mann-Whitney test: *p* = 0.87) or membrane tau (**Fig. 2C**; Mann-Whitney test: *p* = 0.92) of *Fmr1* KO neurons compared with WT. We observed greater input resistance (*R_N_*) in *Fmr1* KO MD neurons compared with WT controls (**Fig. 2D**; Mann-Whitney test: *p* = 0.037, *η*^2^ = 0.052). Based on previous reports of HCN expression in MD neurons (19,27) and extensive literature identifying modifications in HCN expression in *Fmr1* KO mouse models (18,20,22–24,28), we next investigated intrinsic measures of HCN activity to assess potential differences in HCN function between WT and *Fmr1* KO MD neurons.

**Figure 2:**
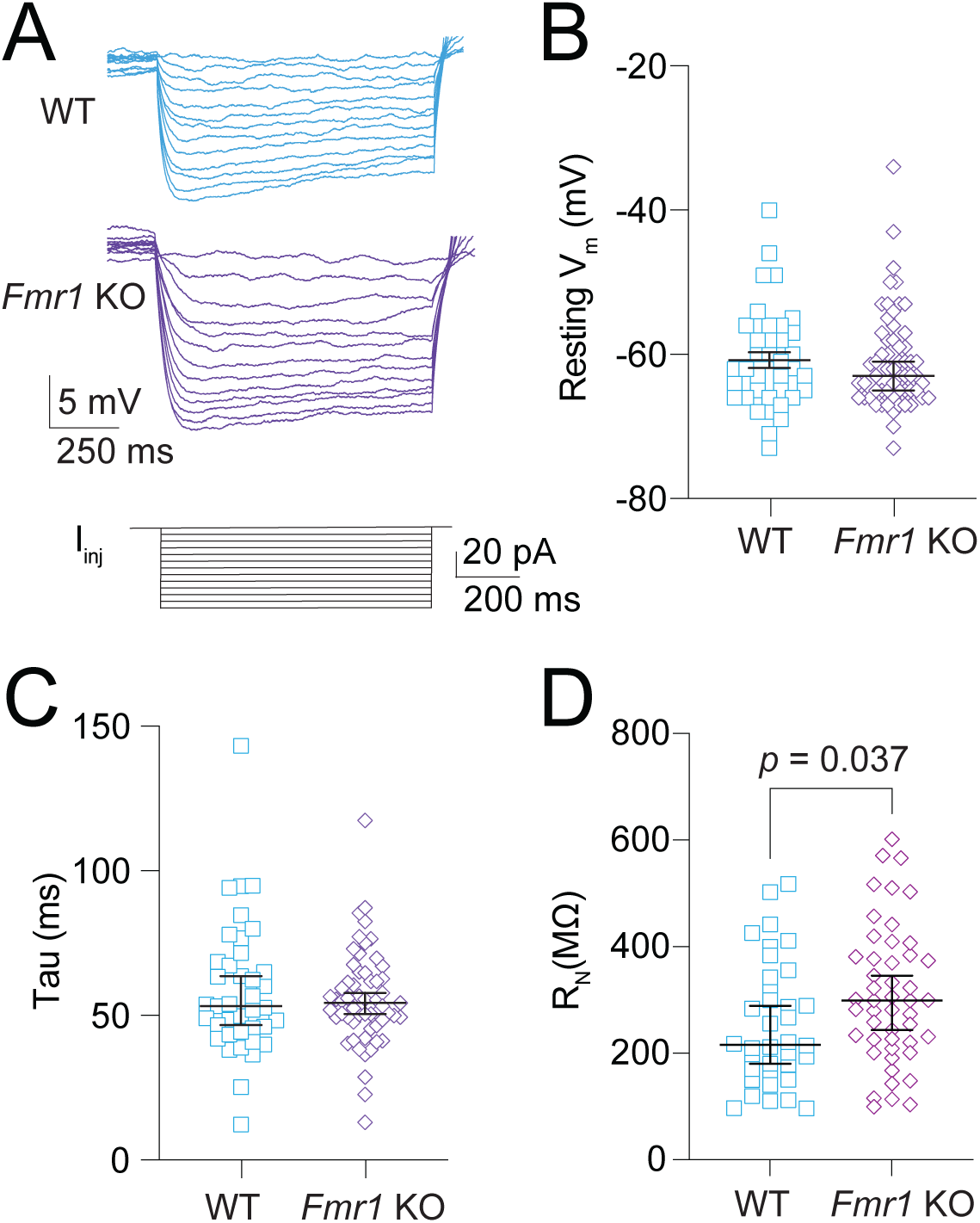
Greater R_N_ in *Fmr1* KO MD neurons compared with WT. **A.** Representative traces showing voltage deflections in response to hyperpolarizing current steps ranging from 0 to -60 pA in 5 pA increments. **B**. Resting V_m_ in WT and *Fmr1* KO neurons. Mann-Whitney test: *p* = 0.87. **C.** Membrane tau of WT and *Fmr1* KO neurons. Mann-Whitney test: *p* = 0.92. **D.** R_N_ in WT and *Fmr1* KO MD neurons measured from the linear portion of each neurons IV plot. Mann-Whitney test: *p* = 0.037.

### Fmr1 KO MD-L neurons have reduced sag when compared with WT MD neurons

We used voltage sag, a well-established measure of HCN activity, to investigate differences in HCN function (**Fig. 3A**). **Figure 3B** shows V-I plots of peak voltage deflection and steady state voltage deflection for both WT and *Fmr1* KO MD-L neurons. While both WT and *Fmr1* KO neurons exhibited clear sag in response to a range of current injections, voltage sag was reduced in *Fmr1* KO neurons compared with WT controls (**Fig. 3C**; 2-way ANOVA: Effect of genotype - *p* = 0.009; *η*^2^ = 0.08). The difference reported in *R_N_* in **Figure 2D** can affect comparisons of sag measured only by current injection. To account for differences in *R_N_* we compared sag in WT and *Fmr1* KO neurons from a peak hyperpolarization of -100 mV. After normalizing for membrane hyperpolarization, the difference in sag persisted (**Fig. 3D**; Mann-Whitney test: *p* = 0.004, *η*^2^ = 0.08). We used the HCN channel blocker ZD7288 (20 μM) to verify that the differences observed in intrinsic neuronal properties stem from HCN channel activity. *R_N_* increased significantly in both WT and *Fmr1* KO neurons with the application of ZD7288 (**Fig. 3E-F**; Wilcoxon test – WT: *p* = 0.002, *η*^2^ = 3.53; *Fmr1* KO: *p* = 0.002, *η*^2^ = 3.14). We additionally measured the effect of ZD7288 on sag. As expected, sag was completely abolished by the wash-on of ZD7288 in both WT and *Fmr1* KO MD neurons (**Fig. 3G-H**; Wilcoxon test – WT: *p* = 0.0005, *η*^2^ = 3.12; *Fmr1* KO: *p* = 0.004, *η*^2^ = 3.16).

**Figure 3:**
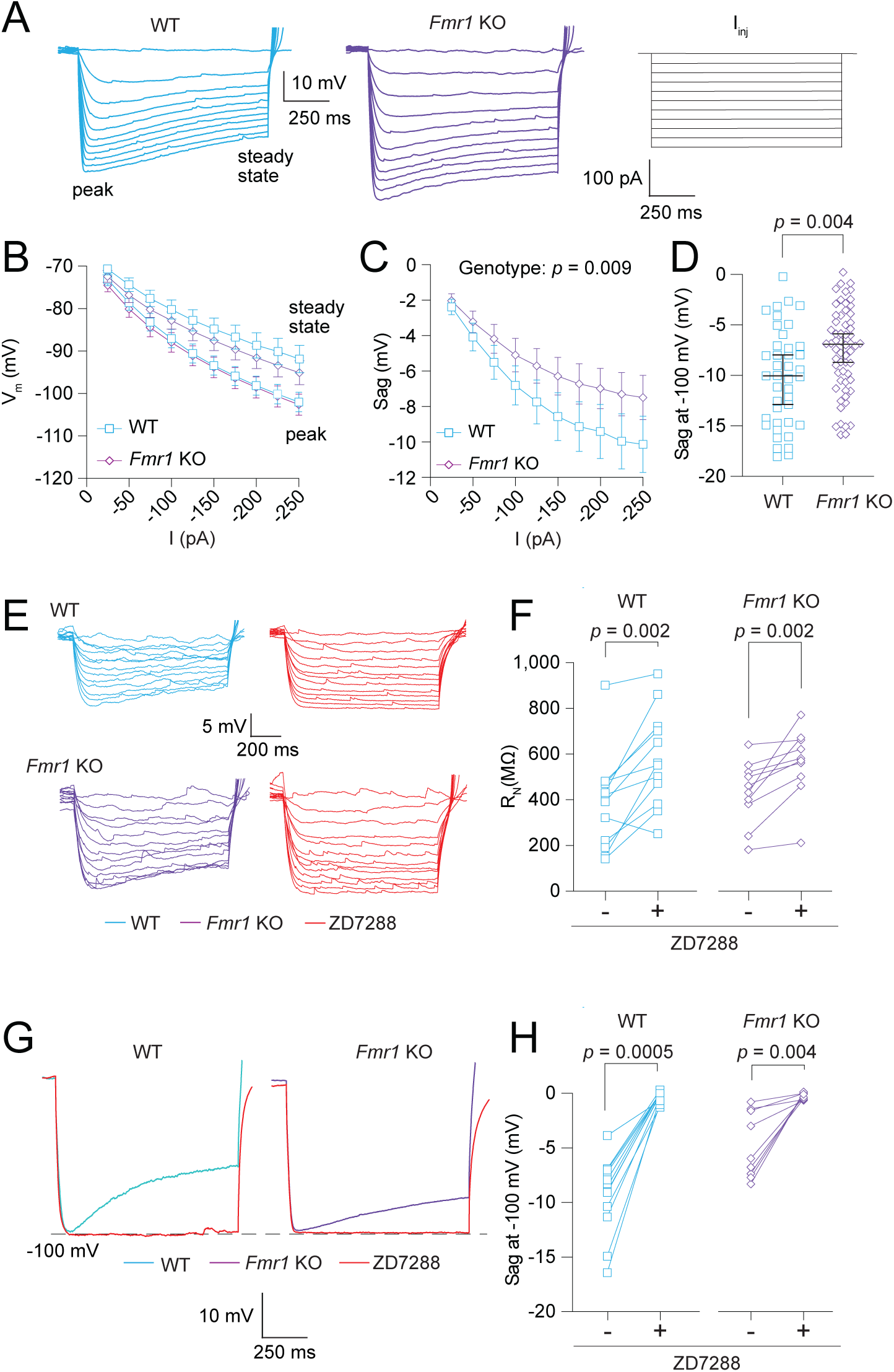
*Fmr1* KO MD neurons exhibit decreased HCN channel activity compared with WT neurons. A. Representative traces of voltage responses from WT and *Fmr1* KO neurons to current stimuli ranging from 0 to -250 pA in 25 pA intervals. **B.** VI plot of WT and *Fmr1* KO neurons measured from both the peak and steady state voltage deflections. **C.** Sag, measured as the difference between peak and steady state voltage deflections, in response to current stimuli from -25 to -250 pA in 25 pA intervals. 2-Way ANOVA – effect of genotype: *p* = 0.009. **D.** Sag in WT and *Fmr1* KO neurons when peak hyperpolarization reached -100 mV. Mann-Whitney test: *p* =0.004. **E.** Representative traces showing WT and *Fmr1* KO neuron voltage responses to current stimuli from 0 to -60 pA in 5 pA intervals before and after the application of 20 μM ZD7288. The linear portion of neuron VI curves were used to calculate R_N_. **F.** Before and after plots of the effect of ZD7288 on WT and *Fmr1* KO R_N_. WT: Wilcoxon test, *p* = 0.002. *Fmr1* KO: Wilcoxon test, *p* = 0.002. **G.** Representative traces showing sag before and after ZD7288 application in WT and *Fmr1* KO neurons. **H.** Before and after plots showing the effect of ZD7288 on sag measurements in WT and *Fmr1* KO neurons. WT: Wilcoxon test: *p* = 0.0005. *Fmr1* KO: Wilcoxon test, *p* = 0.004.

### αEPSP summation is not different between WT and Fmr1 KO MD-L neurons

To test if differences in HCN function between WT and *Fmr1* KO neurons affect signal summation, we injected trains of alpha wave EPSPs (*α*EPSP) at frequencies of 50, 100, and 200 Hz. Stimulus amplitudes were adjusted to approximately 1-2 mV. We found no difference in the amplitude of the first *α*EPSP injected between WT and *Fmr1* KO neurons (Mann-Whitney test: *p* = 0.62). There was no difference in summation between WT and *Fmr1* KO neurons at 50 Hz (**Fig 4A**; 2-way ANOVA – effect of genotype: *p* = 0.23), 100 Hz (**Fig 4B**; 2-way ANOVA – effect of genotype: *p* = 0.54), or 200 Hz (**Fig. 4C**; 2-way ANOVA – effect of genotype: *p* = 0.49). The summation ratio (10^th^ *α*EPSP / 1^st^ *α*EPSP) revealed no difference between WT and *Fmr1* KO neurons (**Fig. 4D**; 2-way ANOVA – effect of genotype: *p* = 0.46). These results suggest that the differences in HCN channel function between WT and *Fmr1* KO neurons do not affect subthreshold summation of *α*EPSP inputs.

**Figure 4:**
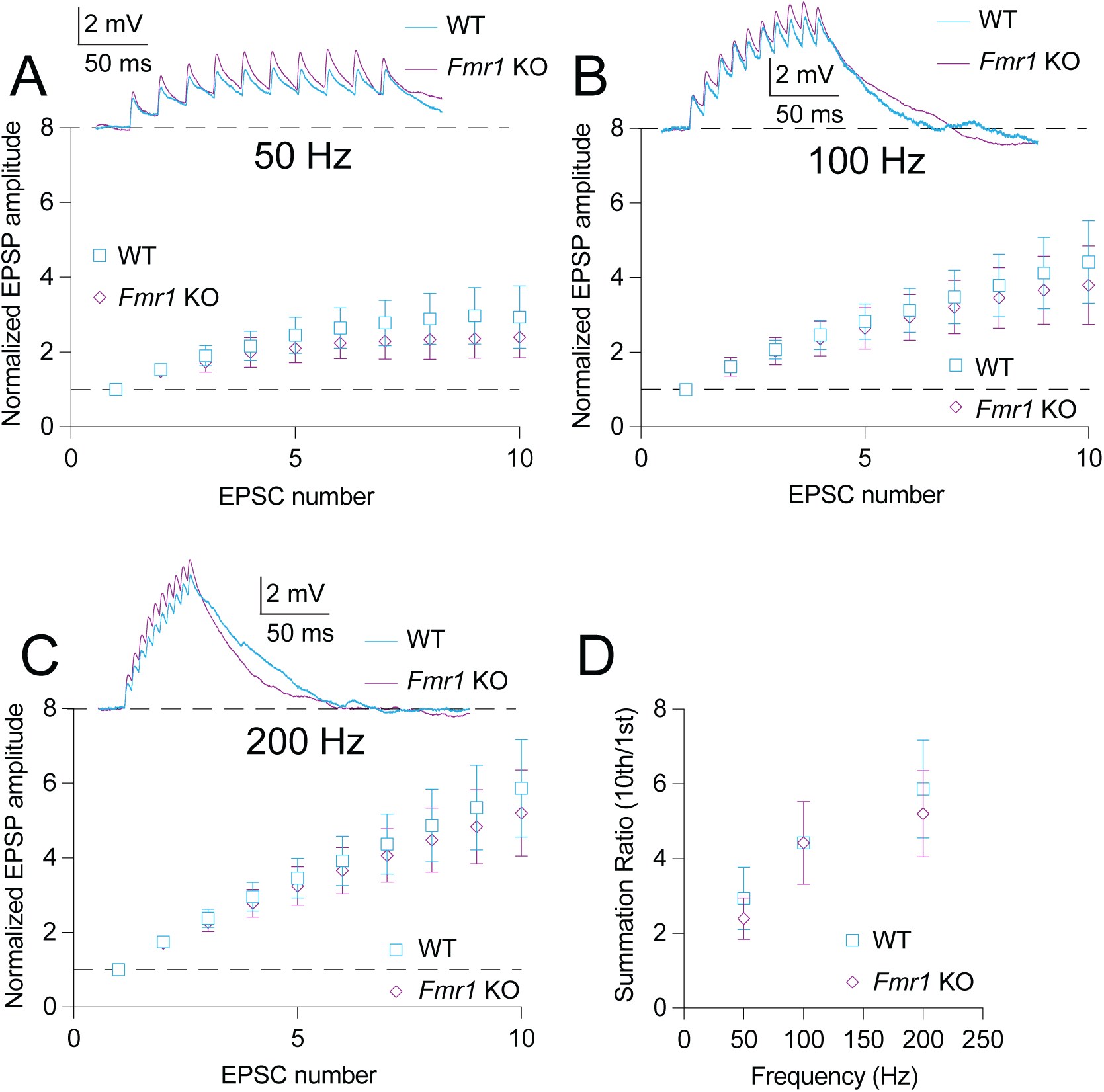
*α*EPSP summation is not different between WT and *Fmr1* KO MD neurons. A. Normalized voltage response to *α*EPSPs delivered at 50 Hz in WT and *Fmr1* KO neurons. 2-Way ANOVA – effect of genotype: *p* = 0.23. **B.** Normalized voltage response to *α*EPSPs delivered at 100 Hz in WT and *Fmr1* KO neurons. 2-Way ANOVA – effect of genotype: *p* = 0.54. **C.** Normalized voltage response to *α*EPSPs delivered at 200 Hz in WT and *Fmr1* KO neurons. 2-Way ANOVA – effect of genotype: *p* = 0.49 **D.** Summation ratio at 50, 100, and 200 Hz *α*EPSP rate in WT and *Fmr1* KO neurons. 2-Way ANOVA – effect of genotype: *p* = 0.46.

### Fmr1 KO MD neurons have greater AHP and delayed RST compared with WT MD neurons

To investigate differences in AHP after a train of action potentials we measured the minimum voltage amplitude reached within 500 ms of the offset of current injection. These measurements were performed both before and after the application of 20 μM ZD7288 (**Fig. 5A**). Measurements of AHP showed a consistent difference in amplitude between WT and *Fmr1* KO neurons (**Fig. 5B**; Mann-Whitney test: *p* = 0.036, *η*^2^ = 0.05). AHP was significantly reduced by the application of ZD7288 in both WT and *Fmr1* KO neurons (**Fig. 5C**; Wilcoxon test – WT: *p* = 0.001, *η*^2^ = 3.13; *Fmr1* KO: *p* = 0.004, *η*^2^ = 3.16). These results suggest that the AHP is strongly influenced by HCN activity and has a reduced amplitude in *Fmr1* KO compared with WT neurons.

**Figure 5:**
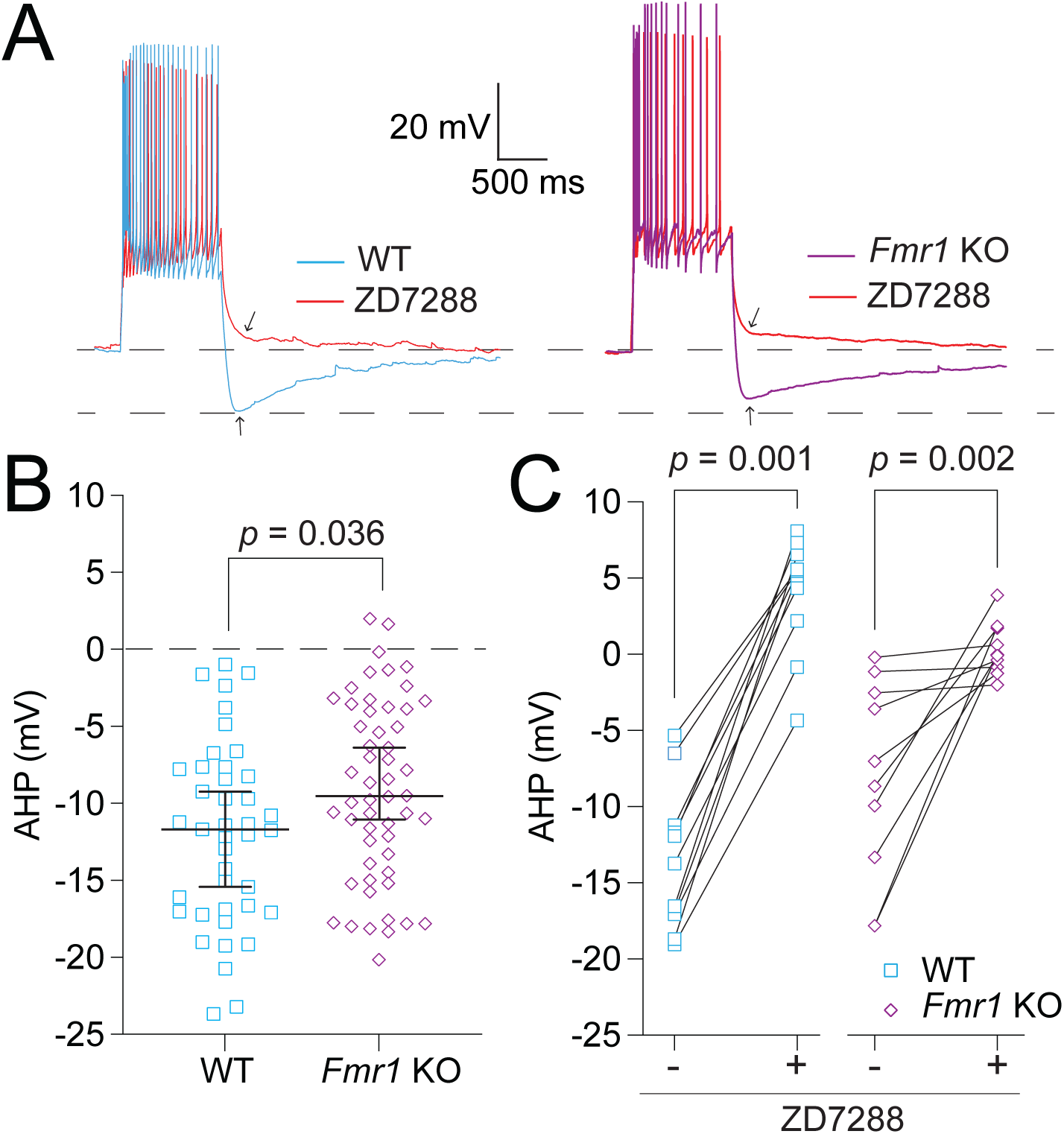
Decreased HCN activity results in decreased AHP amplitude in *Fmr1* KO neurons compared with WT controls. **A.** Representative traces showing AHP amplitude in WT and *Fmr1* KO neurons before and after the application of 20 μM ZD7288. **B.** AHP measured in WT and *Fmr1* KO neurons. Mann-Whitney test: *p* = 0.036. **C.** Before and after plots of the effect of ZD7288 on AHP in WT and *Fmr1* KO neurons. WT: Wilcoxon test, *p* = 0.001. *Fmr1* KO: Wilcoxon test: *p* = 0.002.

We next evaluated RST, defined as the time between the offset of the hyperpolarizing current step and the peak of the first action potential in a rebound burst (**Fig. 6A**). RST measured against hyperpolarizing current step amplitude revealed a delay in RST in *Fmr1* KO neurons compared with WT controls (**Fig. 6B**; Fixed effects (type III) analysis – Effect of genotype: *p* = 0.015, *η*^2^ = 0.07). To account for the peak level of depolarization we plotted RST against peak hyperpolarization for WT and *Fmr1* KO MD neurons (**Fig. 6C**). To further account for level of peak hyperpolarization due to differences in *R_N_* we compared RST when peak hyperpolarization reached -100 mV in WT and *Fmr1* KO neurons and found that *Fmr1* KO neurons show a greater delay in RST compared with WT controls (**Fig. 6D**; Mann-Whitney test: *p* = 0.0014, *η*^2^ = 0.13).

**Figure 6:**
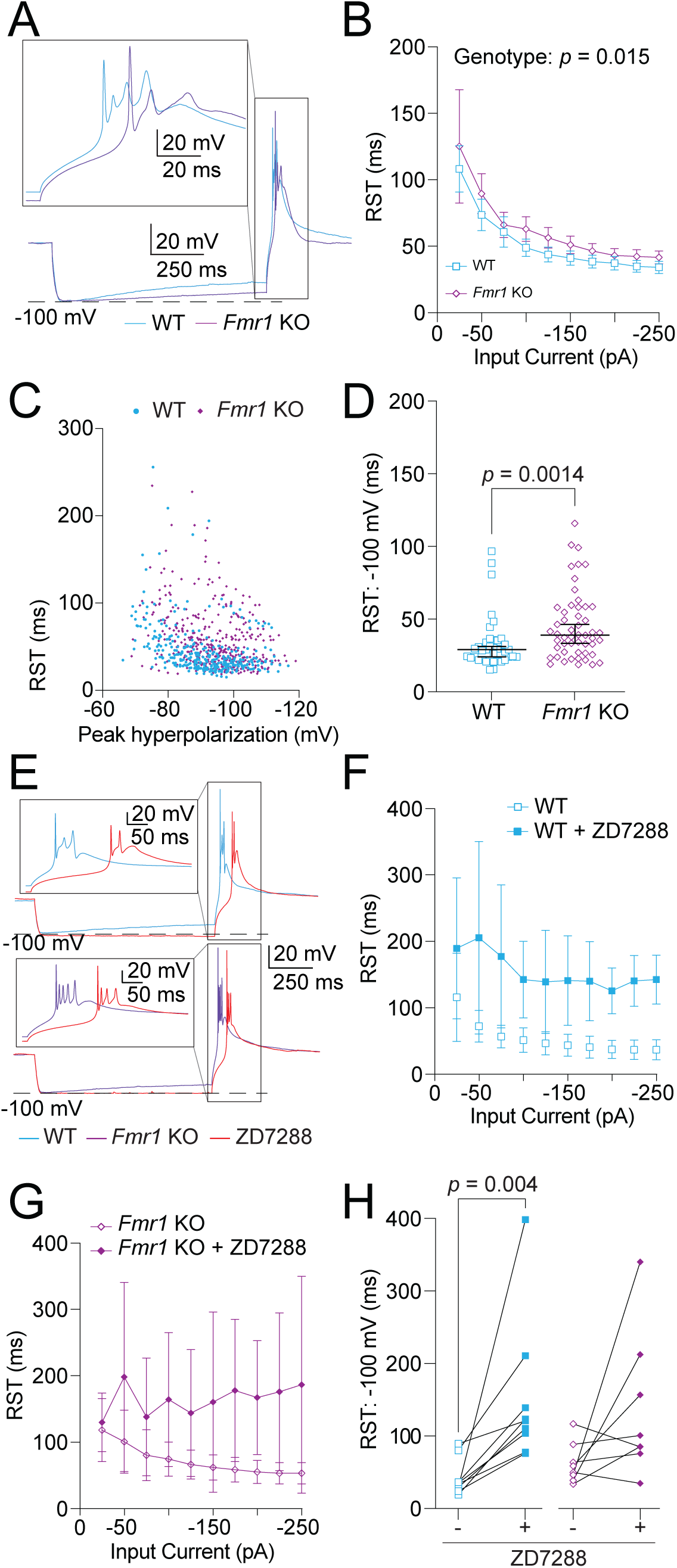
RST is decreased by HCN activity. *Fmr1* KO neurons show a greater delay in RST compared with WT controls. **A.** Representative traces of RST measured as the time from the offset of the hyperpolarizing current to the peak of the first action potential in the burst. Boxed region is expanded in the inset at the top of the panel. **B.** RST measured at current stimuli ranging from 0 to -250 pA in 25 pA intervals. Mixed effects analysis – effect of genotype: *p* = 0.015. **C.** Plot of RST by the peak hyperpolarization reached in response to current stimuli. **D.** RST measured when peak hyperpolarization reached -100 mV. Mann-Whitney test: *p* = 0.0014. **E.** Representative traces showing rebound bursts before and after the application of 20 μM ZD7288 in WT and *Fmr1* KO neurons. Insets represent expanded views of the boxed regions. **F.** RST measurements in response to hyperpolarizing current stimuli from 0 to -250 pA in 25 pA intervals in WT MD neurons before and after wash-on of 20 μM ZD7288. WT Fixed effects (type III) analysis – effect of genotype: *p* = 0.015. **G.** RST measurements in response to hyperpolarizing current stimuli from 0 to -250 pA in 25 pA intervals in *Fmr1* KO MD neurons before and after wash-on of 20 μM ZD7288. *Fmr1* KO Fixed effects (type III) analysis – effect of genotype: *p* = 0.032. **H.** Before and after plots of RST showing the effects of 20 μM ZD7288 wash-on in WT and *Fmr1* KO neurons measured when peak hyperpolarization reached -100 mV. WT: Wilcoxon test, *p* = 0.004. *Fmr1* KO: Wilcoxon test, *p* = 0.56.

In the absence of rebound burst firing, the release of hyperpolarization causes an HCN channel-mediated depolarization before returning to rest (29) . We hypothesized that this effect contributes to the difference in RST observed here. To test HCN involvement, we measured RST before and after wash-on of 20 μM ZD7288 (**Fig. 6E**). When compared against current amplitude both WT (**Fig. 6F**; WT Fixed effects (type III) analysis – effect of genotype: *p* = 0.015, *η*^2^ = 0.42) and *Fmr1* KO (**Fig 6G**; *Fmr1* KO Fixed effects (type III) analysis – effect of genotype: *p* = 0.032, *η*^2^ = 0.23) neurons showed a significant delay in RST with blockade of HCN channels. When normalized to steps in which peak hyperpolarization reached -100 mV, only WT neurons showed a significant difference in RST after ZD7288 wash-on (**Fig. 6H**; Wilcoxon test – WT: *p* = 0.016, *η*^2^ = 3.2; *Fmr1* KO: *p* = 0.56), suggesting that decreased HCN channel activity results in delayed RST in *Fmr1* KO neurons.

### Burst firing and calcium spike properties are not different between WT and Fmr1 KO MD neurons

Thalamic neurons display state-dependent firing properties based on the availability of Ca_v_3 channels: neurons fire tonic spikes when resting at depolarized potentials (when Ca_V_3 channels are inactivated) and fire bursts when hyperpolarized (when Ca_V_3 channels are available) (30). **Figs. 7-8** investigate differences in action potential firing based on neuron state and stimulus type. Tonic firing in WT and *Fmr1* KO neurons was investigated and no differences were identified (**Supp.** Fig. 1).

**Figure 7:**
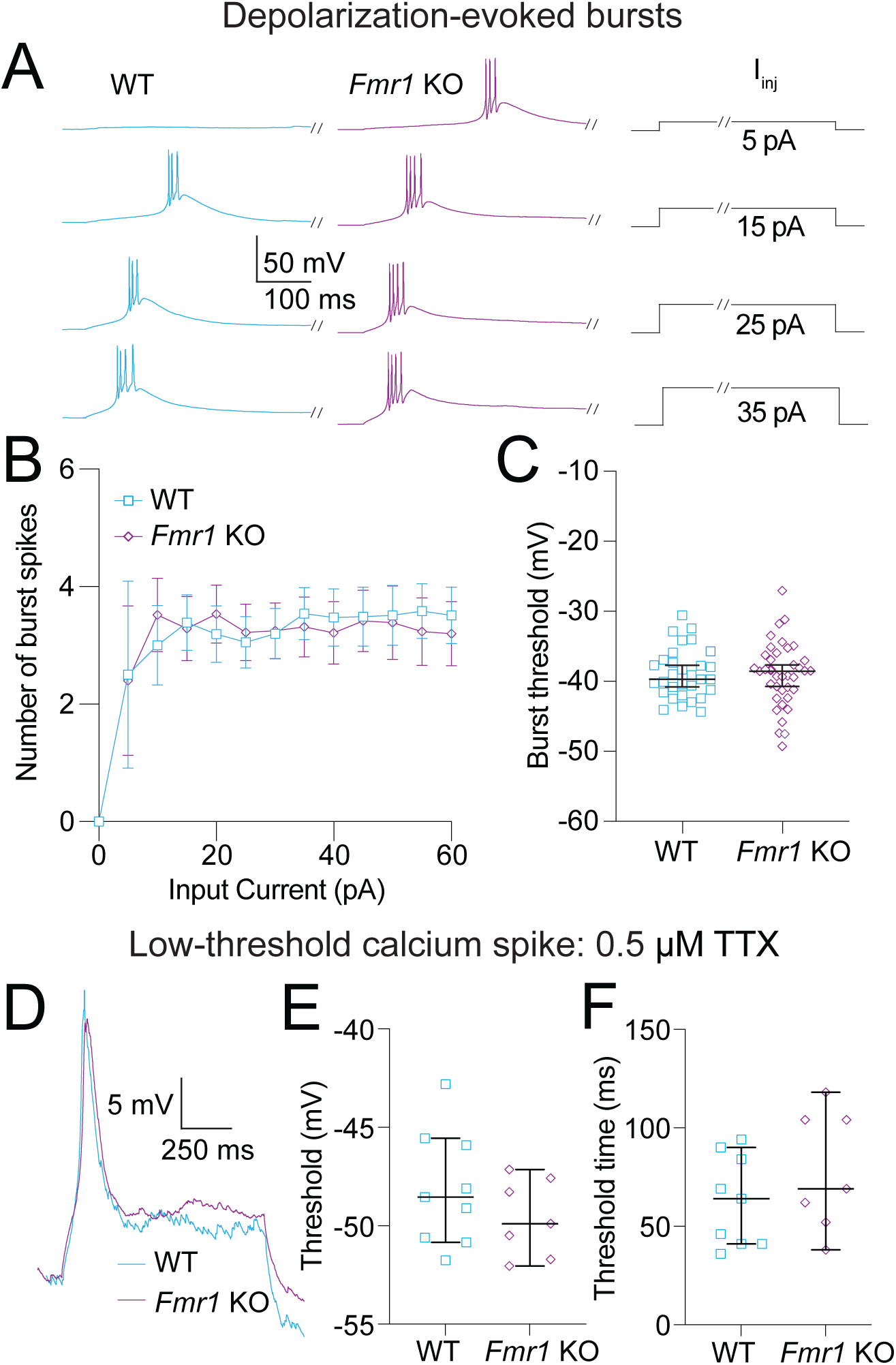
Bursting evoked by depolarizing current steps is not different between WT and *Fmr1* KO MD neurons. **A.** Representative traces showing burst firing in response to 5, 15, 25, and 35 pA steps. Slashes on current steps show where voltage traces were truncated to better show bursts. **B.** Number of action potentials per burst for current steps from 0 to 60 pA in 5 pA intervals. Fixed effects (type III) analysis – effect of genotype: *p* = 0.51. **C.** Burst threshold measured from the first step to elicit a burst. Mann-Whitney test: *p* = 0.81. **D.** Representative traces of LTS isolated by the application of 0.5 μM TTX. **E.** LTS threshold measured from the first current step to evoke a spike. Mann-Whitney test: *p* = 0.47. **F.** LTS time to threshold measured from the first current step to evoke a spike. Mann-Whitney test: *p* = 0.31.

We recorded bursts of action potentials in response to depolarizing current steps ranging from 0 to 60 pA in 5 pA intervals (**Fig. 7A**). There were no differences in either the number of action potentials fired in each burst (**Fig. 7B**; Fixed effects (type III) analysis – effect of genotype: *p* = 0.51) or the threshold for the first action potential in the first step to elicit a burst (**Fig. 7C**; Mann-Whitney test: *p* = 0.81). To compare calcium spikes evoked by depolarization we applied the sodium channel blocker TTX (0.5 μM) (**Fig. 7D**). We found no difference in LTS threshold (**Fig. 7E**; Mann-Whitney test: *p* = 0.47), or the time from the offset of the current injection and the peak of the LTS (Mann-Whitney test: *p* = 0.47) between WT and *Fmr1* KO neurons. Added variability from the slower activation time of Ca_v_3 channels compared with Na_v_ channels may obscure differences in LTS onset. To address this confound we measured the latency from the onset of current to LTS threshold. We found no difference in spike timing using this measure (**Fig. 7F**; Mann-Whitney: *p* = 0.31).

We then recorded the properties of rebound bursts evoked in response to hyperpolarizing current steps from 0 to -60 pA in 5 pA intervals (**Fig. 8A**). There was no difference between WT and *Fmr1* KO neurons in the number of action potentials fired in each rebound burst (**Fig. 8B**; Fixed effects (type III) analysis – effect of genotype: *p* = 0.9) or the rebound burst threshold (**Fig. 8C**; Mann-Whitney test: *p* = 0.37). With the addition of TTX to isolate calcium spikes (**Fig. 8D**) we found no difference in spike threshold (**Fig. 8E**; Mann-Whitney test: *p* = 0.87) or the timing between current offset and spike peak (Mann-Whitney test: *p* = 0.2) between WT and *Fmr1* KO neurons. As in **Fig 7F**, we measured the time to LTS threshold to assess LTS latency using more concise measure. We found that the latency to LTS threshold was greater in *Fmr1* KO compared with WT MD neurons after the offset of hyperpolarizing current (**Fig. 8F**; Mann-Whitney: *p* = 0.028, *η*^2^ = 0.22). Our results show waveform properties of rebound bursts and LTS associated with Ca_v_3 channels are not different between *Fmr1* KO and WT MD neurons. Consistent with measurements of RST in **Fig. 6**, we identified greater latency to hyperpolarization-evoked rebound LTS in *Fmr1* KO compared with WT MD neurons.

**Figure 8:**
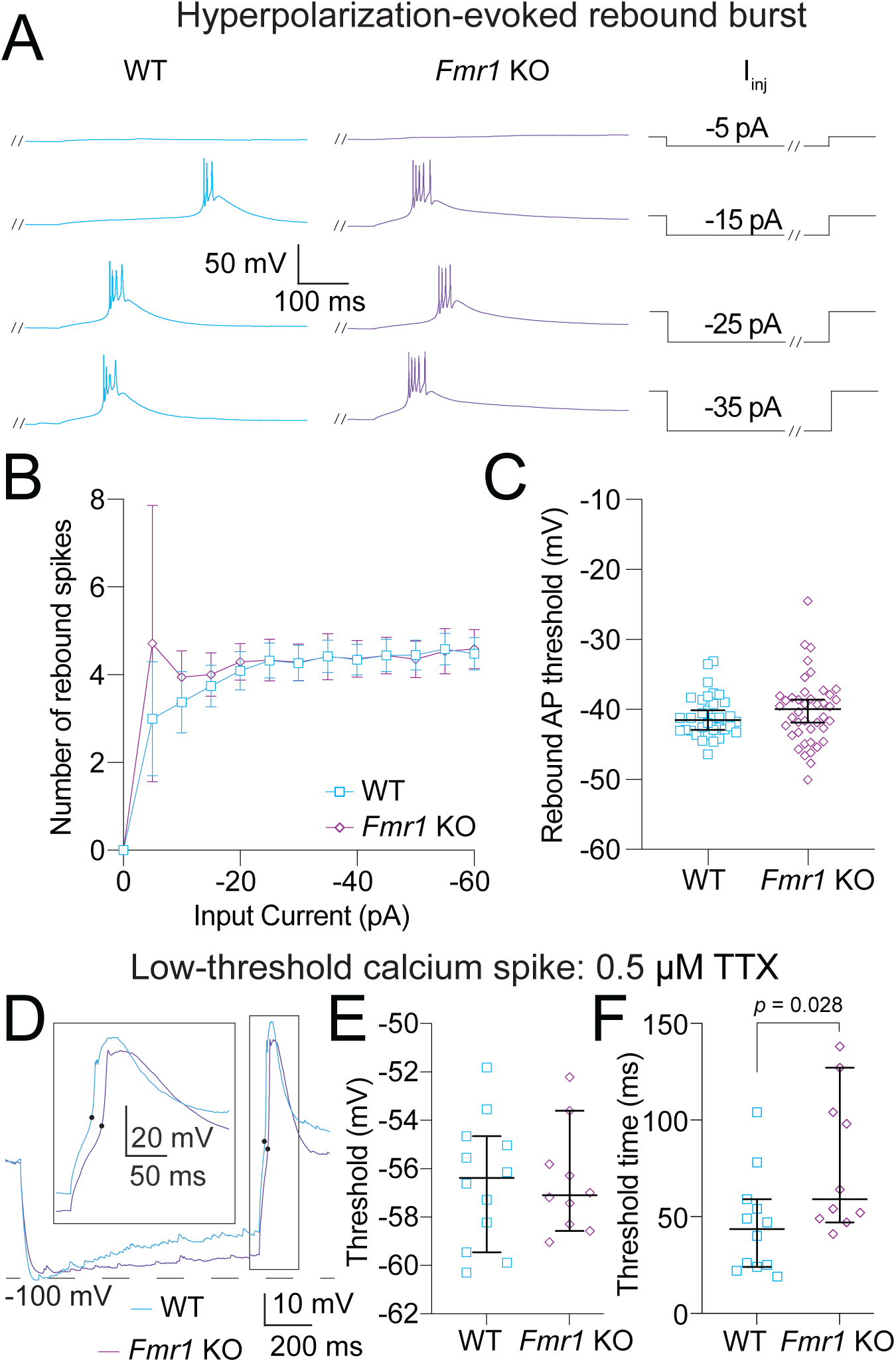
Bursting evoked by hyperpolarizing current steps is not different between WT and *Fmr1* KO MD neurons. **A.** Representative traces showing burst firing in response to -5, - 15, -25, and -35 pA steps. Slashes on current steps show where voltage traces were truncated to better show bursts. **B.** Number of action potentials per burst for current steps from 0 to -60 pA in 5 pA intervals. Fixed effects (type III) analysis – effect of genotype: *p* = 0.9. **C.** Burst threshold measured from the first hyperpolarizing step to elicit a burst. Mann-Whitney test: *p* = 0.37. **D.** Representative traces of LTSs isolated by the application of 0.5 μM TTX. **E.** LTS threshold measured from the first hyperpolarizing current step to evoke a spike. Mann-Whitney test: *p* = 0.87. Black dots represent threshold measurements for LTS. **F.** LTS time to threshold measured from the first hyperpolarizing current step to evoke a spike. Mann-Whitney test: *p* = 0.028.

## Discussion

Reciprocal connectivity between MD and mPFC is required for higher order operations like executive function and social behavior (2,3,6,31) . The symptomatology of FXS displays a high degree of overlap with functions controlled by thalamocortical circuitry (32,33), providing strong implications of prefrontal circuit pathology in FXS. Research has revealed disruptions to prefrontal circuitry through changes in both intrinsic and synaptic properties as well as in mPFC-mediated learning (18,34,35). The importance of MD in executive and affective functions led to our hypothesis that disruptions in MD function may contribute to pathology associated with FXS. We found that *Fmr1* KO MD-L neurons that project to mPFC displayed reduced HCN activity compared with WT MD-L neurons. Further investigation revealed HCN-channel-dependent differences in membrane voltage dynamics after the offset of depolarizing and hyperpolarizing current injections manifesting as decreased AHP amplitude and delayed RST.

We identified differences in events following the offset of both hyperpolarizing (RST) and depolarizing (AHP) current stimuli when comparing *Fmr1* KO with WT MD-L neurons. AHP amplitude was decreased, and RST was delayed in bursts and LTS in *Fmr1* KO neurons when compared to WT controls. These findings suggest that *Fmr1* KO MD-L neurons may have difficulty maintaining phase locked rhythmic firing, which is critical for normal thalamocortical circuit function. The thalamus is important for the generation and maintenance of rhythmic activity in the brain (30). Furthermore, rhythmic activity in MD is an integral component to normal higher order brain function, particularly working memory (36,37) and normal sleep function (38,39).

Because *Fmr1* KO MD-L neurons had increased input resistance, we predicted that modeled synaptic currents would show greater summation in *Fmr1* KO compared with WT neurons. However, injected *α*EPSPs revealed no difference in summation properties at any recorded frequency. This contrasts with observed effects of HCN channel activity on EPSP summation in hippocampal pyramidal neurons (40). It remains unknown how HCN channels localize within the membrane of MD neurons. Dendritic HCN channel localization may impact *α*EPSPs differences observed in somatic recordings (41). Although this study focuses on differences in HCN channel activity, it is likely that other channelopathies are present in MD neurons of *Fmr1* KO animals (42). For instance, the lack of difference in *α*EPSP summation between *Fmr1* KO neurons could be explained by increased resting K^+^ conductance in *Fmr1* KO MD-L neurons. MD neurons express 2-pore K^+^ channels (K_2P_) which contribute to the control over state-dependent firing within thalamic neurons (43). Increased activity of these channels could account for the lack of difference in *α*EPSP summation between WT and *Fmr1* KO MD-L neurons despite the difference in input resistance reported here.

Recordings from human MD reveal the presence of both sleep spindles and ripples, establishing a role for MD in normal sleep function (44). Spindle activity in thalamic neurons is thought to stem from rebound bursting in response to hyperpolarizing GABAergic inputs (45). Reduced HCN activity in *Fmr1* KO MD-L neurons contributes to delayed spike timing and reduced AHP amplitude, changes that will functionally impair a neuron’s ability to generate rhythmic bursting activity. Rhythmic bursting in thalamic neurons depends on HCN channel activity as is shown by disrupted burst generation in an *Hcn4* conditional KO mouse (46). Sleep is disrupted in patients with FXS (47,48). *Fmr1* KO mice have reduced sleep spindle density and sleep spindle discoordination in cortical areas including the prefrontal network (49). Activation of G-protein receptor coupled inward rectifying K^+^ (GIRK) channels restored normal cortical sleep spindle activity in *Fmr1* KO mice (50). Our results implicate disruption of rhythmic bursting activity in MD neurons in *Fmr1* KO mice that are likely involved in sleep disruptions observed in both mice models of FXS and humans with FXS.

In this study we performed whole cell current clamp recordings from CTB-labeled MD neurons that project to mPFC in WT and *Fmr1* KO MD-M and MD-L neurons. This study focused on identifying the effects of *Fmr1* KO on intrinsic neuronal properties in MD, however, synaptic dysfunction is widely reported in FXS (51) . HCN plays a prominent role in the integration, filtering, and coordination of synaptic inputs in neuronal dendrites (40,52–54). Although somatically-generated *α*EPSPs did not reveal differences in subthreshold summation between WT and *Fmr1* KO MD-L neurons, the localization and dendritic effects of HCN channel expression remain unexplored. Thalamic neurons, through their unique ion channel expression profile, generate and maintain rhythmic activity. Coupled with the widespread interconnectivity between cortical and subcortical brain structures, MD is a promising target for therapeutic intervention for the treatment of symptoms associated with FXS.

## ACKNOWLEDGEMENTS

We thank Meredith McCarty, Aurora Weiden, Madelynn Campbell, Joy Adler, and Mendee Geist for their technical assistance. We thank members of the Brumback and Howard labs (especially Jessica Chancey and MacKenzie Howard), Darrin Brager, and Daniel Johnston for helpful discussions.

## CONTRIBUTIONS

G.J.O. wrote the paper and interpreted and analyzed data. P.L. performed electrophysiological recordings. A.K. performed imaging and projection tracing. A.B. conceived of the project, designed all experiments, wrote all code, analyzed data, and wrote portions of the paper. All authors approved the final manuscript prior to publication.

## DATA AVAILABILITY/ SUPPLEMENTAL MATERIALS

Source data and supplemental materials are available through github.

## References

1. Fuster J, Alexander G (1973): Firing changes in cells of the nucleus medialis dorsalis associated with delayed response behavior. Brain Res 61: 79–91.

2. Parnaudeau S, O’Neill P-K, Bolkan SS, Ward RD, Abbas AI, Roth BL, et al. (2013): Inhibition of Mediodorsal Thalamus Disrupts Thalamofrontal Connectivity and Cognition. Neuron 77: 1151–1162.

3. Bolkan SS, Stujenske JM, Parnaudeau S, Spellman TJ, Rauffenbart C, Abbas AI, et al. (2017): Thalamic projections sustain prefrontal activity during working memory maintenance. Nature Neuroscience 20: 987–996.

4. Alexander GE, Fuster JM (1973): Effects of cooling prefrontal cortex on cell firing in the nucleus medialis dorsalis. Brain Res 61: 93–105.

5. Ouhaz Z, Fleming H, Mitchell AS (2018): Cognitive Functions and Neurodevelopmental Disorders Involving the Prefrontal Cortex and Mediodorsal Thalamus. Front Neurosci 12: 33.

6. Rikhye RV, Gilra A, Halassa MM (2018): Thalamic regulation of switching between cortical representations enables cognitive flexibility. Nat Neurosci 21: 1753–1763.

7. Lee S, Ahmed T, Lee S, Kim H, Choi S, Kim D-S, et al. (2011): Bidirectional modulation of fear extinction by mediodorsal thalamic firing in mice. Nat Neurosci 15: 308–314.

8. Hwang K, Bruss J, Tranel D, Boes AD (2020): Network Localization of Executive Function Deficits in Patients with Focal Thalamic Lesions. J Cognitive Neurosci 28: 1–16.

9. Hwang K, Shine JM, Cole MW, Sorenson E (2022): Thalamocortical contribution to cognitive task activity. Elife 11. 10.7554/elife.81282

10. Kloet S de, Bruinsma B, Terra H, Heistek T, Passchier E, Berg A van den, et al. (2020): Bi-directional command of cognitive control by distinct prefrontal cortical output neurons to thalamus and striatum. Nat Commun 12: 1994.

11. Nair A, Treiber JM, Shukla DK, Shih P, Müller R-A (2013): Impaired thalamocortical connectivity in autism spectrum disorder: a study of functional and anatomical connectivity. Brain 136: 1942–1955.

12. Woodward ND, Giraldo-Chica M, Rogers B, Cascio CJ (2017): Thalamocortical dysconnectivity in autism spectrum disorder: An analysis of the Autism Brain Imaging Data Exchange. Biological psychiatry Cognitive neuroscience and neuroimaging 2: 76–84.

13. Lanfranchi S, Cornoldi C, Drigo S, Vianello R (2009): Working memory in individuals with fragile X syndrome. Child neuropsychology : a journal on normal and abnormal development in childhood and adolescence 15: 105 119.

14. D’Hooge R, Nagels G, Franck F, Bakker CE, Reyniers E, Storm K, et al. (1997): Mildly impaired water maze performance in maleFmr1 knockout mice. Neuroscience 76: 367–376.

15. Kooy RF, D’Hooge R, Reyniers E, Bakker CE, Nagels G, Boulle KD, et al. (1996): Transgenic mouse model for the fragile X syndrome. Am J Méd Genet 64: 241–245.

16. Mercaldo V, Vidimova B, Gastaldo D, Fernández E, Lo AC, Cencelli G, et al. (2023): Altered striatal actin dynamics drives behavioral inflexibility in a mouse model of fragile X syndrome. Neuron. 10.1016/j.neuron.2023.03.008

17. Brumback AC, Ellwood IT, Kjaerby C, Iafrati J, Robinson S, Lee AT, et al. (2017): Identifying specific prefrontal neurons that contribute to autism-associated abnormalities in physiology and social behavior. Mol Psychiatr 23: 2078–2089.

18. Kalmbach BE, Johnston D, Brager DH (2015): Cell-Type Specific Channelopathies in the Prefrontal Cortex of the fmr1-/y Mouse Model of Fragile X Syndrome. Eneuro 2: ENEURO.0114-15.2015.

19. Lyuboslavsky P, Ordemann GJ, Kizimenko A, Brumback AC (2024): Two contrasting mediodorsal thalamic circuits target the mouse medial prefrontal cortex. J Neurophysiol. 10.1152/jn.00456.2023

20. Brager DH, Akhavan AR, Johnston D (2012): Impaired Dendritic Expression and Plasticity of h-Channels in the fmr1−/y Mouse Model of Fragile X Syndrome. Cell Reports 1: 225–233.

21. Ordemann GJ, Apgar CJ, Chitwood RA, Brager DH (2021): Altered A-type potassium channel function impairs dendritic spike initiation and temporoammonic long-term potentiation in Fragile X syndrome. J Neurosci JN-RM-0082–21.

22. Deng P-Y, Klyachko VA (2016): Genetic upregulation of BK channel activity normalizes multiple synaptic and circuit defects in a mouse model of fragile X syndrome. The Journal of Physiology 594: 83–97.

23. Deng P-Y, Klyachko VA (2016): Increased Persistent Sodium Current Causes Neuronal Hyperexcitability in the Entorhinal Cortex of Fmr1 Knockout Mice. Cell Reports 16: 3157–3166.

24. Brandalise F, Kalmbach BE, Mehta P, Thornton O, Johnston D, Zemelman BV, Brager DH (2020): Fragile X Mental Retardation Protein Bidirectionally Controls Dendritic I h in a Cell Type-Specific Manner between Mouse Hippocampus and Prefrontal Cortex. J Neurosci 40: 5327–5340.

25. Mátyás F, Lee J, Shin H, Acsády L (2014): The fear circuit of the mouse forebrain: connections between the mediodorsal thalamus, frontal cortices and basolateral amygdala. European Journal of Neuroscience 39: 1810–1823.

26. Cohen J (1988): Statistical Power Analysis for the Behavioral Sciences. 10.4324/9780203771587

27. Notomi T, Shigemoto R (2004): Immunohistochemical localization of Ih channel subunits, HCN1–4, in the rat brain. J Comp Neurol 471: 241–276.

28. Deng P-Y, Avraham O, Cavalli V, Klyachko VA (2021): Hyperexcitability of Sensory Neurons in Fragile X Mouse Model. Front Mol Neurosci 14: 796053.

29. Mishra P, Narayanan R (2023): The enigmatic HCN channels: A cellular neurophysiology perspective. Proteins: Struct, Funct, Bioinform. 10.1002/prot.26643

30. Llinás RR, Steriade M (2006): Bursting of Thalamic Neurons and States of Vigilance. J Neurophysiol 95: 3297–3308.

31. Pergola G, Danet L, Pitel A-L, Carlesimo GA, Segobin S, Pariente J, et al. (2018): The Regulatory Role of the Human Mediodorsal Thalamus. Trends in Cognitive Sciences. 10.1016/j.tics.2018.08.006

32. Schmitt LM, Li J, Liu R, Horn PS, Sweeney JA, Erickson CA, Pedapati EV (2022): Altered frontal connectivity as a mechanism for executive function deficits in fragile X syndrome. Mol Autism 13: 47.

33. Holsen LM, Dalton KM, Johnstone T, Davidson RJ (2008): Prefrontal social cognition network dysfunction underlying face encoding and social anxiety in fragile X syndrome. NeuroImage 43: 592–604.

34. Krueger DD, Osterweil EK, Chen SP, Tye LD, Bear MF (2011): Cognitive dysfunction and prefrontal synaptic abnormalities in a mouse model of fragile X syndrome. Proc National Acad Sci 108: 2587–2592.

35. Siegel JJ, Chitwood RA, Ding JM, Payne C, Taylor W, Gray R, et al. (2017): Prefrontal Cortex Dysfunction in Fragile X Mice Depends on the Continued Absence of Fragile X Mental Retardation Protein in the Adult Brain. The Journal of Neuroscience 37: 7305–7317.

36. Parnaudeau S, Bolkan SS, Kellendonk C (2017): The Mediodorsal Thalamus: An Essential Partner of the Prefrontal Cortex for Cognition. Biol Psychiat 83: 648–656.

37. Rikhye RV, Wimmer RD, Halassa MM (2018): Toward an Integrative Theory of Thalamic Function. Annu Rev Neurosci 41: 1–21.

38. Halgren M, Ulbert I, Bastuji H, Fabó D, Erőss L, Rey M, et al. (2019): The generation and propagation of the human alpha rhythm. Proc Natl Acad Sci 116: 23772–23782.

39. Mak-McCully RA, Rolland M, Sargsyan A, Gonzalez C, Magnin M, Chauvel P, et al. (2017): Coordination of cortical and thalamic activity during non-REM sleep in humans. Nat Commun 8: 15499.

40. Poolos NP, Migliore M, Johnston D (2002): Pharmacological upregulation of h-channels reduces the excitability of pyramidal neuron dendrites. Nat Neurosci 5: 767–774.

41. Johnston D, Wu SM-S (1994): Foundations of Cellular Neurophysiology. The MIT Press.

42. Brager DH, Johnston D (2014): Channelopathies and dendritic dysfunction in fragile X syndrome. Brain Res Bull 103: 11–17.

43. Bista P, Pawlowski M, Cerina M, Ehling P, Leist M, Meuth P, et al. (2015): Differential phospholipase C-dependent modulation of TASK and TREK two-pore domain K+ channels in rat thalamocortical relay neurons. J Physiology 593: 127–144.

44. Szalárdy O, Simor P, Ujma PP, Jordán Z, Halász L, Erőss L, et al. (2024): Temporal association between sleep spindles and ripples in the human anterior and mediodorsal thalamus. Eur J Neurosci 59: 641–661.

45. Steriade M (2006): Grouping of brain rhythms in corticothalamic systems. Neuroscience 137: 1087–1106.

46. Zobeiri M, Chaudhary R, Blaich A, Rottmann M, Herrmann S, Meuth P, et al. (2019): The Hyperpolarization-Activated HCN4 Channel is Important for Proper Maintenance of Oscillatory Activity in the Thalamocortical System. Cereb Cortex bhz047-.

47. Budimirovic DB, Protic DD, Delahunty CM, Andrews HF, Choo T, Xu Q, et al. (2022): Sleep problems in fragile X syndrome: Cross-sectional analysis of a large clinic-based cohort. Am J Méd Genet Part A 188: 1029–1039.

48. Kronk R, Bishop EE, Raspa M, Bickel JO, Mandel DA, Bailey DB (2010): Prevalence, Nature, and Correlates of Sleep Problems Among Children with Fragile X Syndrome Based on a Large Scale Parent Survey. Sleep 33: 679–687.

49. Saré RM, Harkless L, Levine M, Torossian A, Sheeler CA, Smith CB (2017): Deficient Sleep in Mouse Models of Fragile X Syndrome. Frontiers in Molecular Neuroscience 10: 280.

50. Martinez JD, Wilson LG, Brancaleone WP, Peterson KG, Popke DS, Garzon VC, et al. (2024): Hypnotic treatment improves sleep architecture and EEG disruptions and rescues memory deficits in a mouse model of fragile X syndrome. Cell Rep 43: 114266.

51. Bagni C, Zukin RS (2019): A Synaptic Perspective of Fragile X Syndrome and Autism Spectrum Disorders. Neuron 101: 1070–1088.

52. Magee JC, Cook EP (2000): Somatic EPSP amplitude is independent of synapse location in hippocampal pyramidal neurons. Nat Neurosci 3: 895–903.

53. Narayanan R, Johnston D (2007): Long-Term Potentiation in Rat Hippocampal Neurons Is Accompanied by Spatially Widespread Changes in Intrinsic Oscillatory Dynamics and Excitability. Neuron 56: 1061–1075.

54. Berger T, Senn W, Lüscher H-R (2003): Hyperpolarization-Activated Current Ih Disconnects Somatic and Dendritic Spike Initiation Zones in Layer V Pyramidal Neurons. J Neurophysiol 90: 2428–2437.

