## Supplemental Data for "*Fmr1* KO causes delayed rebound spike timing in mediodorsal thalamocortical neurons through regulation of HCN channel activity"

Tonic firing: Depolarized resting membrane potential

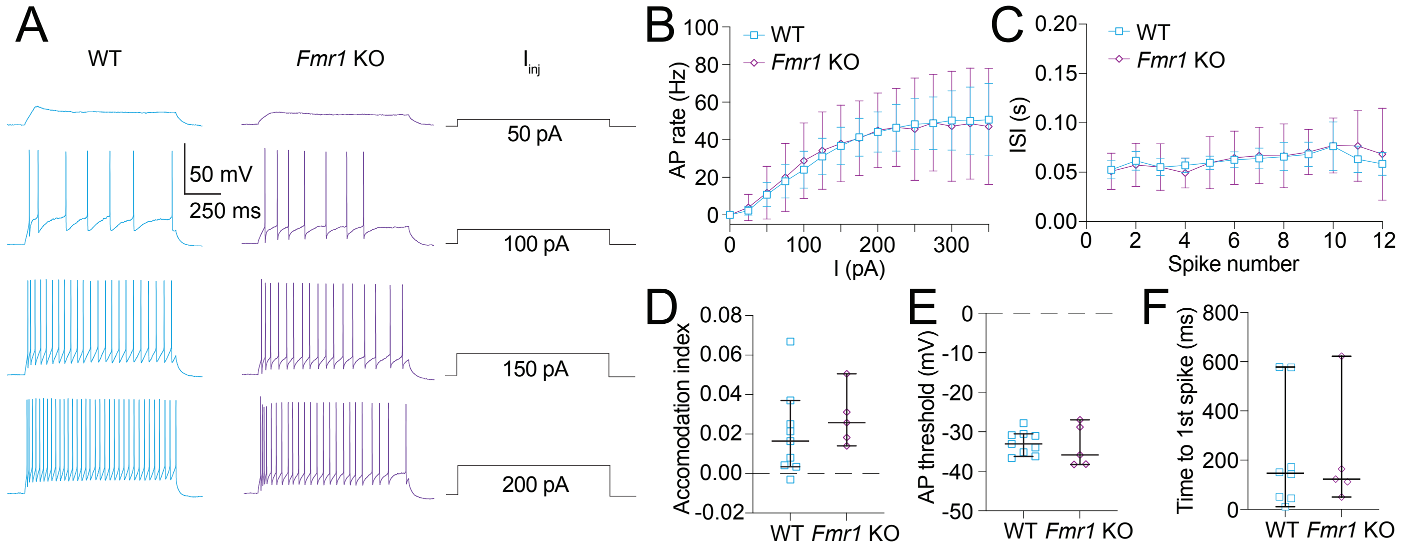


Tonic firing: 20 μM Mibefradil

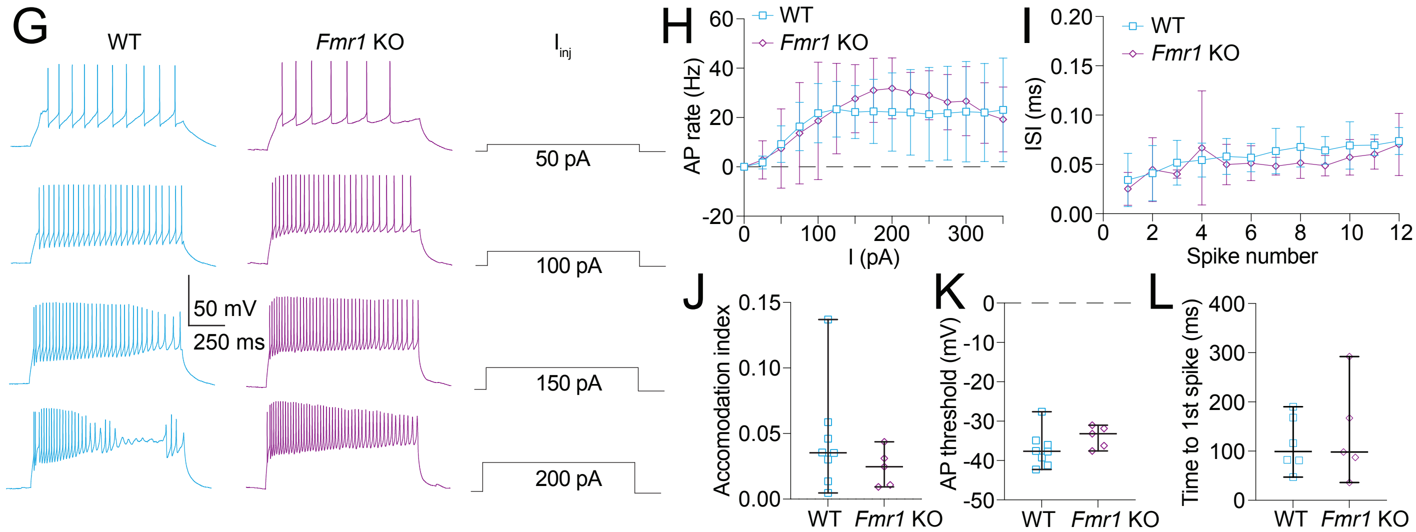


**Figure S1: Tonic action potential firing is not different between WT and *Fmr1* KO MD neurons. A-F.** Tonic firing data collected from cells that did not fire bursts at resting membrane potential. **A.** Representative traces showing tonic action potential firing in WT and *Fmr1* KO neurons in response to 50, 100, 150, and 200 pA current steps. **B.** Action potential firing rate measured at current steps from 0 to 350 in 25 pA intervals. 2-way ANOVA – effect of genotype: $p$ = 0.97. **C.** ISI of action potentials in the first step with 12 or more action potentials. Fixed effects (type III) – effect of genotype: $p$ = 0.93. **D.** Accomodation index measurements taken from the first current step to elicit 12 or more action potentials. Mann-Whitney: $p$ = 0.3. **E.** Action potential threshold measured from the first step with 12 or more action potentials. Mann-Whitney: $p$ = 0.7. **F.** Time to the peak of the first spike in the first current step to elicit one or more action potentials. Mann-Whitney: $p$ = 0.94. **G-L.** Tonic firing data collected from cells held at -65 mV with the CaV 3 blocker mibefradil added to prevent bursting. **G.** Representative traces showing tonic action potential firing in WT and *Fmr1* KO neurons in response to 50, 100, 150, and 200 pA current steps. **H.** Action potential firing rate measured at current steps from 0 to 350 in 25 pA intervals. 2-way ANOVA – effect of genotype: $p$ = 0.67. **I.** ISI of action potentials in the first step with 12 or more action potentials. Fixed effects (type III) analysis – effect of genotype: $p$ = 0.26. **J.** Accomodation index measurements taken from the first current step to elicit 12 or more action potentials. Mann-Whitney test: $p$ = 0.28. **K.** Action potential threshold measured from the first step with 12 or more action potentials. Mann-Whitney test: $p$ = 0.14. **L.** Time to the peak of the first spike in the first current step to elicit one or more action potentials. Mann-Whitney test: $p$ = 0.93.

***Tonic action potential firing is not different between WT and Fmr1 KO MD neurons***
To isolate tonic action potential firing, we took two approaches. First, we analyzed that did not fire bursts at RMP (**Fig. S1A-F**). Second, we held cells at -65 mV while blocking $Ca_v3$ channels with 20 μM mibefradil, which prevents burst firing (**Fig. S1G-L**). We used measures of firing frequency, ISI, spike threshold, and accommodation to assess differences in tonic firing between WT and *Fmr1* KO MD neurons.

No comparison of tonic firing properties at RMP was different between WT and *Fmr1* KO neurons (**Fig S1A-F**). We measured action potential firing rate (**Fig S1B**; 2-way ANOVA – effect of genotype: $p = 0.97$), ISI (**Fig. S1C**; Fixed effects (type III) – effect of genotype: $p = 0.93$), accommodation (**Fig. S1D**; Mann-Whitney test: $p = 0.3$), action potential threshold (**Fig. S1E**; Mann-Whitney test: $p = 0.7$), and time to first action potential in the first depolarizing step to elicit an action potential (**Fig. S1F**; Mann-Whitney test: $p = 0.94$) and found no differences.

Similarly, no comparison of tonic firing properties during mibefradil wash-on was different between WT and *Fmr1* KO neurons (**Fig S1G-L**). No difference was identified in measurements of action potential frequency (**Fig S1H**; 2-way ANOVA – effect of genotype: $p = 0.67$), ISI (**Fig. S1I**; Fixed effects (type III) analysis – effect of genotype: $p = 0.26$), accommodation (**Fig. S1J**; Mann-Whitney test: $p = 0.28$), action potential threshold (**Fig. S1K**; Mann-Whitney test: $p = 0.14$), or the time to first action potential in the first current step to elicit an action potential (**Fig. S1L**; Mann-Whitney test: $p = 0.93$). These results show that suprathreshold tonic firing is not affected in MD neurons in *Fmr1* KO animals.

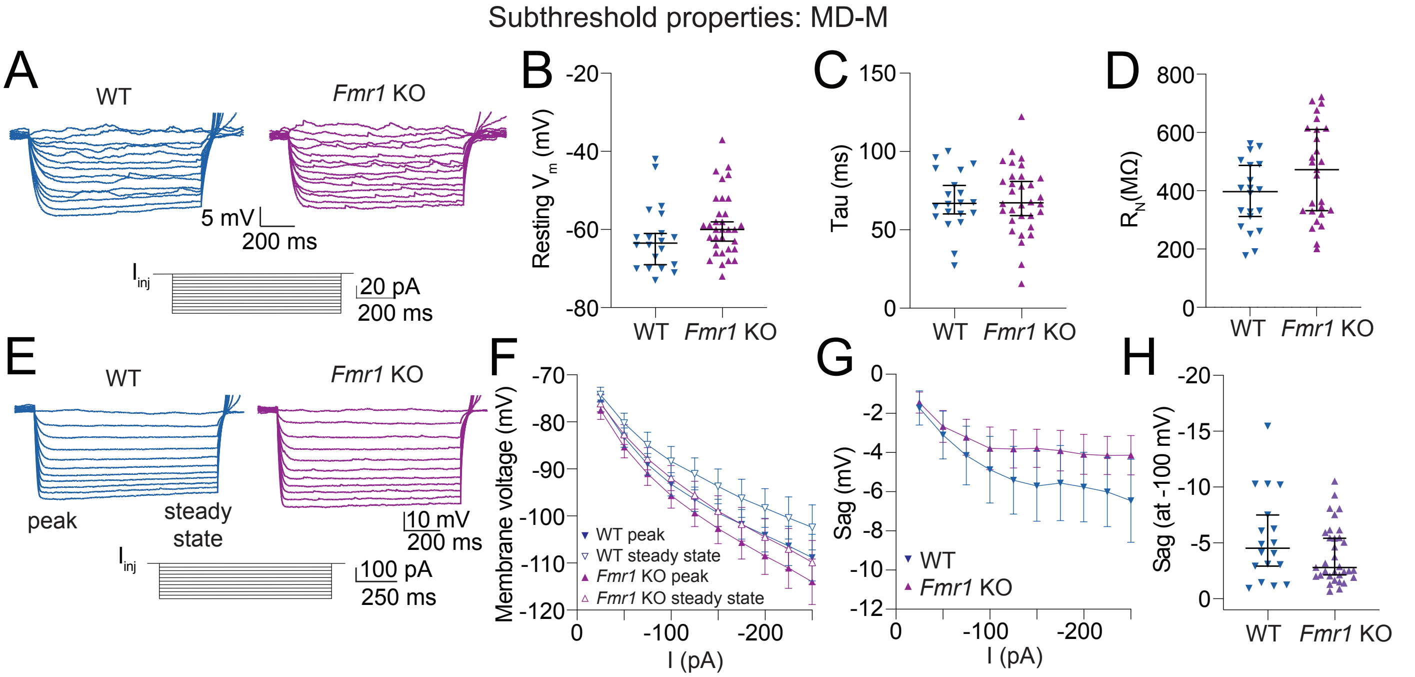


**Figure S2: WT and *Fmr1* KO MD-M neurons show no difference in subthreshold properties. A.** Representative traces showing voltage responses to hyperpolarizing current steps from 0 to -60 pA in 5 pA intervals in WT and *Fmr1* KO neurons. **B.** Resting membrane potential in WT and *Fmr1* KO neurons. Mann-Whitney test: $p$ = 0.09. **C.** Membrane tau measured from a step to -10 pA in WT and *Fmr1* KO neurons. Mann-Whitney test: $p$ = 0.86. **D.** Input resistance measurements from the linear portion of the IV for WT and *Fmr1* KO MD-M neurons. Mann-Whitney test: $p$ = 0.09. **E.** Representative traces of voltage responses from WT and *Fmr1* KO MD-M neurons to current stimuli ranging from 0 to -250 pA in 25 pA intervals. **F.** VI plot of WT and *Fmr1* KO neurons measured from both the peak and steady state voltage deflections. **G.** Sag, measured as the difference between peak and steady state voltage deflections, in response to current stimuli from -25 to -250 pA in 25 pA intervals. 2-way ANOVA – effect of genotype: $p$ = 0.083. **H.** Sag in WT and *Fmr1* KO neurons when peak hyperpolarization reached -100 mV. Mann-Whitney test: $p$ = 0.24.

***Subthreshold properties are not different between WT and Fmr1 KO MD-M neurons***

All data discussed so far were from the lateral subnucleus of MD projecting to mPFC (MD-L→mPFC neurons). We tested if these same findings were present in neurons in the medial subnucleus of MD (MD-M→mPFC neurons). We measured voltage responses to current stimuli from -60 to 60 pA in 5 pA intervals (**Fig. S2A**) in WT and *Fmr1* KO MD-M neurons. We found no differences in measurements of resting membrane potential (**Fig. S2B**; Mann-Whitney test: $p$ = 0.09), membrane tau (**Fig. S2C**; Mann-Whitney test: $p$ = 0.86), or $R_N$ (**Fig. S2D**; Mann-Whitney test: $p$ = 0.09). We further investigated the subthreshold property of sag to assess whether changes in HCN activity observed in MD-L were also present in MD-M. We recorded voltage responses ranging from -250 to 0 pA in 25 pA intervals in WT and *Fmr1* KO MD-M neurons (**Fig. S2E**). We created a VI plot from the peak and steady state portions of voltage traces in response to each current step for both WT and *Fmr1* KO MD-M neurons (**Fig. S2F**) to calculate voltage sag at each current step. There was no difference in sag between WT and *Fmr1* KO MD-M neurons either when measured at each current injection amplitude (**Fig. S2G**; 2-way ANOVA – effect of genotype: $p$ = 0.083) or when traces with a peak hyperpolarization of -100 mV were analyzed (**Fig. S2H**; Mann-Whitney test: $p$ = 0.24).

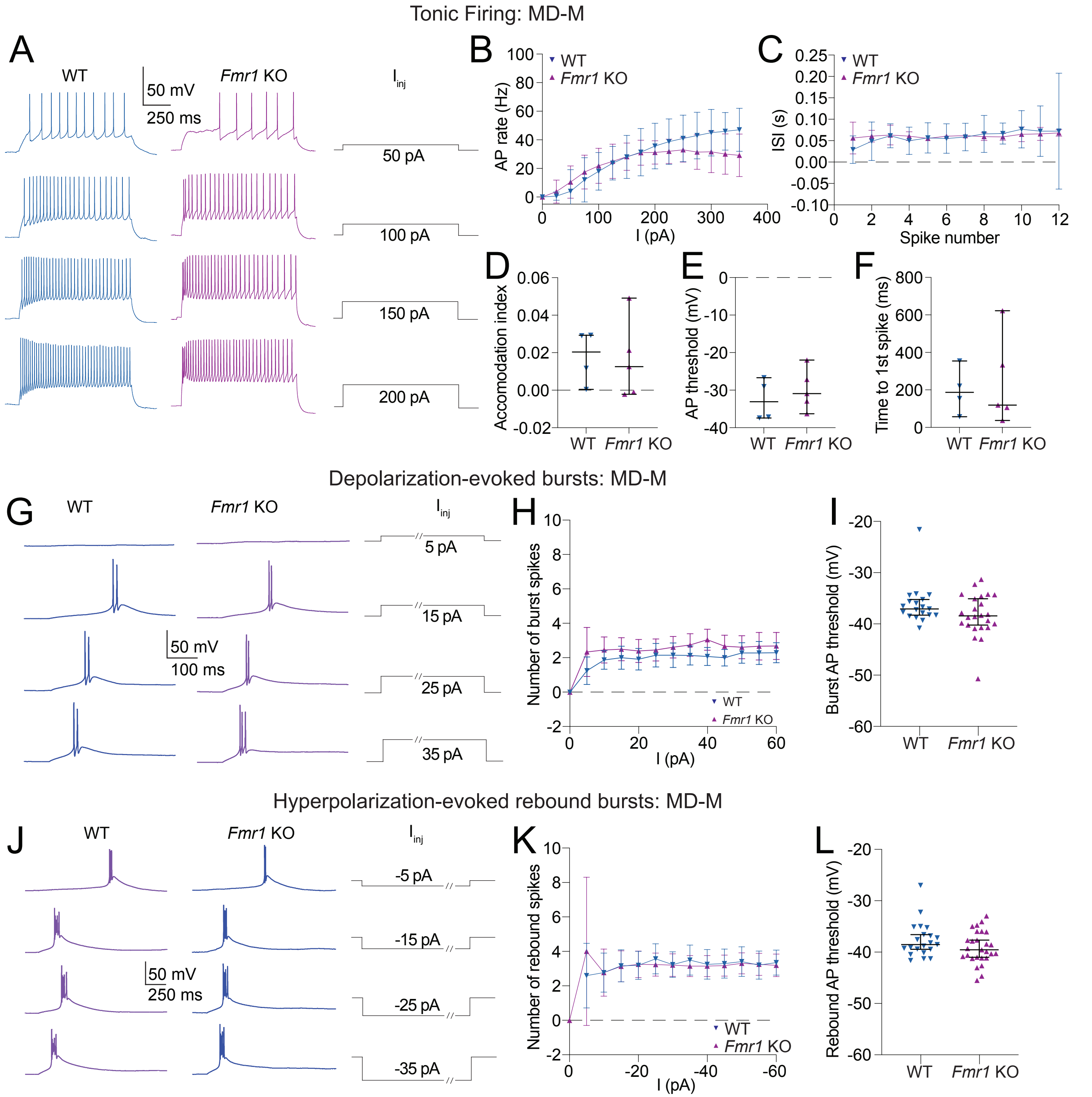


**Figure S3: WT and *Fmr1* KO MD-M neurons show no difference in suprathreshold properties. A-F.** Tonic firing data collected from cells that did not fire bursts at resting membrane potential. **A.** Representative traces showing tonic action potential firing in WT and *Fmr1* KO MD-M neurons in response to 50, 100, 150, and 200 pA current steps. **B.** Action potential firing rate measured at current steps from 0 to 350 in 25 pA intervals. 2-way ANOVA – effect of genotype: $p = 0.35$. **C.** ISI of action potentials in the first step with 12 or more action potentials. Fixed effects (type III) analysis – effect of genotype: $p = 0.7$. **D.** Accomodation index measurements taken from the first current step to elicit 12 or more action potentials. Mann-Whitney test: $p = 0.73$. **E.** Action potential threshold measured from the first step with 12 or more action potentials. Mann-Whitney test: $p = 0.56$. **F.** Time to the peak of the first spike in the first current step to elicit any action potentials. Mann-Whitney test: $p = 0.9$. **G.** Representative traces showing burst firing in response to 5, 15, 25, and 35 pA steps. Slashes on current steps show where voltage traces were truncated to better visualize bursts. **H.** Number of action potentials per burst for current steps from 0 to 60 pA in 5 pA intervals. Fixed effects (type III) analysis – effect of genotype: $p = 0.32$. **I.** Burst threshold measured from the first step to elicit a burst. Mann-Whitney test: $p = 0.16$. **J.** Representative traces showing burst firing in response to -5, -15, -25, and -35 pA steps. Slashes on current steps show where voltage traces were truncated to better show bursts. **K.** Number of action potentials per burst for current steps from 0 to -60 pA in 5 pA intervals. Fixed effects (type III) analysis – effect of genotype: $p = 0.9$. **L.** Burst threshold measured from the first hyperpolarizing step to elicit a burst. Mann-Whitney test: $p = 0.18$.

***Suprathreshold properties are not different between WT and Fmr1 KO MD-M neurons***

We investigated tonic firing from WT and *Fmr1* KO MD-M neurons that did not fire bursts at resting membrane potential (**Fig. S3A**). Our experiments revealed no differences in action potential firing (**Fig. S3B**; 2-way ANOVA – effect of genotype: $p = 0.35$) or ISI (**Fig. S3C**; Fixed effects (type III) analysis – effect of genotype: $p = 0.7$). We found no differences in properties associated with changes in voltage gated channels associated with suprathreshold activity in WT and *Fmr1* KO MD-M neurons including accommodation (**Fig. S3D**; Mann-Whitney test: $p = 0.73$), action potential threshold (**Fig. S3E**; Mann-Whitney test: $p = 0.56$), and time to first action potential in the first step to elicit action potentials (**Fig. S3F**; Mann-Whitney test: $p = 0.9$).

Burst firing was analyzed in depolarizing current steps from 0 to 60 pA in 5 pA intervals (**Fig. S3G**). No difference in either the number of action potentials per burst (**Fig. S3H**; Fixed effects (type III) analysis – effect of genotype: $p = 0.32$) or in the burst action potential threshold (**Fig. S3I**; Mann-Whitney test: $p = 0.16$) was identified between WT and *Fmr1* KO MD-M neurons.

Rebound burst firing was analyzed after offset of hyperpolarizing current steps from -60 to 0 pA in 5 pA intervals (**Fig. S3J**). No difference in either the number of action potentials per burst (**Fig. S3K**; Fixed effects (type III) analysis – effect of genotype: $p = 0.9$) or in the rebound burst action potential threshold (**Fig. S3L**; Mann-Whitney test: $p = 0.18$) was identified between WT and *Fmr1* KO MD-M neurons.

| Resource Type | Specific Reagent or Resource | Source or Reference | Identifiers |
|---|---|---|---|
| Add additional rows as needed for each resource type | Include species and sex when applicable. | Include name of manufacturer, company, repository, individual, or research lab. | Include catalog numbers, stock numbers, database IDs or accession numbers, and/or RRIDs. RRIDs are highly encouraged. |
| Bacterial or Viral Strain | Cholera toxin subunit B (CTB) | Molecular Probes - Thermo Fisher Scientific | Cat# C22841 / RRID: |
| Chemical Compound or Drug | tetrodotoxin citrate | Abcam | Cat# ab120055 / RRID: |
| Chemical Compound or Drug | ZD7288 | Hello Bio | Cat# HB1152 / RRID: |
| Organism/Strain | Mouse: C57Bl6 | Jackson Laboratory | Stock #000664 / RRID:MGI:3028467 |
| Organism/Strain | Mouse: Fmr1 -/y | Kimberley Huber, Ph.D. (Jackson Laboratory) | Stock#003025 / RRID:IMSR_JAX:003025 |
| Software; Algorithm | Clampex | Axon Intruments | RRID:SCR_011323 |
| Software; Algorithm | Matlab | Wavemetrics | RRID:SCR_001622 |
| Software; Algorithm | Prism | GraphPad | RRID:SCR_002798 |
| Software; Algorithm | Adobe Illustrator | Adobe | RRID:SCR_010279 |
| Chemical Compound or Drug | Mibefradil | Tocris | Cat#2198 |
